# CRISP enables comparisons of image-based spatial transcriptomic segmentation quality across ten organs

**DOI:** 10.64898/2026.04.16.718947

**Authors:** James R. Rose, Elex S. Rose, José A. F. Assumpção, Heather Pathak, Hannah E. Peck, Loren E. Sasser, Charmy J. Patel, Daryll Vanover, Philip J. Santangelo

## Abstract

Image-based spatial transcriptomics depends on cell segmentation to assign transcripts to individual cells, but how segmentation algorithms perform across tissues with distinct cellular architectures is poorly understood. This study presents the broadest independent benchmark to date of cell segmentation algorithms for spatial transcriptomics, comparing five approaches across ten mouse tissues using a 5,006-gene Xenium panel. To quantify segmentation errors, Co-expression Rejection in Segmentation Purity (CRISP) was developed, an open-source tool available in R and Python that measures cell purity through tissue-specific mutually exclusive marker co-expression without requiring ground truth annotations. This benchmark revealed that segmentation algorithms face a fundamental tradeoff between maximizing transcript capture and maintaining cell purity, and that the severity of this tradeoff is tissue-dependent. Proseg achieved the highest average performance across tissues, though the magnitude of its advantage varies with tissue architecture. Overall, CRISP provides per-tissue performance profiles as a practical resource for algorithm selection.

## Introduction

While methods for spatially resolving gene expression have been in development since the late 1980s^1^, recent years have marked an inflection point in the adoption of spatially-resolved transcriptomics (SRT) driven by commercially available image-based platforms that have made subcellular transcript detection accessible to a broader research community^2,3^. SRT aims to quantify and locate expressed RNA molecules in biological tissues in real physical space, and image-based methods tend to excel at both tasks on the transcript level. These methods, however, are also dependent on accurate cell segmentation to correctly assign transcripts to cells and enable downstream tasks such as cell typing, clustering, and differential expression testing^4,5^. As a result, the biological utility of these platforms is often limited not only by their ability to resolve and detect transcripts but also by how faithfully those transcripts are assigned to the correct cell, making segmentation arguably the most consequential computational step in the image-based SRT workflow.

Methods developed to segment cells in the context of SRT can be broadly organized by the primary data modality they leverage. Image-based approaches use microscopy and dye or immuno-based staining to identify and highlight cell boundaries. Cellpose is a popular example used to perform cell segmentation primarily on non-SRT image data and has also been used in SRT applications, particularly when paired with staining techniques such as H&E or nucleus staining with DAPI^6^. Early approaches in SRT often relied on algorithms like Cellpose for nuclear expansion, in which DAPI-stained nuclei were segmented from images and uniformly expanded at a given diameter to approximate cell boundaries. While this expansion increases the number of transcripts captured per cell, it comes at the cost of increased noise from inappropriate cell segmentations, particularly in tissues with large cells or cells with non-uniform morphology^7^. More recently, image-based methods, such as the 10x multimodal segmentation algorithm, pair nuclear staining with additional immunofluorescence channels and machine learning models in order to more accurately delineate cell boundaries.

Transcript-based segmentation algorithms use a fundamentally different strategy, relying on the spatial coordinates of detected RNA molecules rather than image data to infer cell boundaries. Baysor was an early method developed to utilize the shorter distances between neighboring transcripts within cells compared to extracellular regions in order to perform cell segmentation^8^. Proseg similarly works on transcript location data but instead conducts voxel-level simulations to iteratively fit cell boundaries to observed transcript distributions^9^. While primarily transcript-based, both methods can optionally incorporate image-derived nuclear or other segmentations as priors. Newer methods have explored graph-based methods, including Segger, which constructs heterogeneous graphs of neighboring transcripts and nuclei, applying graph neural networks to assign transcripts to cells^10^. These categories carry practical implications: image-based methods are constrained by staining quality and morphological assumptions, while transcript-based methods can in principle adapt to any tissue architecture but may be sensitive to transcript density and spatial noise.

The effects of inaccurate segmentation have been shown to significantly confound downstream analysis of SRT data^4^, yet researchers currently lack comprehensive guidance for selecting among available algorithms. Existing benchmarks have primarily compared across platforms rather than across segmentation approaches within a single platform or tissue. Ren et al. evaluated multiple SRT technologies across three cancer tissue types, finding that the Xenium platform outperformed other commercial offerings, while Wang et al. compared platforms within human tissue microarrays and observed evidence of inappropriate co-expression of mutually exclusive marker genes in default Xenium-segmented data^11,12^. While these benchmarks are important, neither study systematically varied the segmentation algorithm applied to the same underlying data. Efforts to directly quantify segmentation quality have introduced useful metrics, most notably the mutually exclusive co-expression rate (MECR) developed by Hartman & Satija^13^. MECR, however, lacks an available software implementation and was evaluated only in mouse brain tissue. More recently, SpatialQM extended this purity-based logic across six tissues alongside other quality control metrics, describing a negative relationship between transcript capture and purity as well as improved purity using Proseg^14^. However, the authors did not discuss tissue-specific differences in this tradeoff, nor did they benchmark more than two segmentation algorithms. To the authors’ knowledge, no independent study has yet compared more than two segmentation algorithms in image-based SRT data across a broad range of tissue types, leaving a critical gap in understanding how algorithm choice interacts with unique cell morphology of different tissues to shape data quality.

To address these limitations, Co-expression Rejection in Segmentation Purity (CRISP; https://github.com/santangelo-lab/CRISP) was developed, a software package available in both R and Python that builds on the MECR concept and extends it into a practical, tissue-aware framework for quantifying segmentation purity through detection of inappropriate co-expression of mutually exclusive marker genes. CRISP was applied alongside a comprehensive suite of metrics to benchmark five segmentation approaches across a Xenium 5K panel tissue microarray (TMA) dataset spanning 10 mouse tissue types. These approaches include nuclear segmentation with no expansion, the 10x multimodal segmentation kit, Baysor (with both nuclear and 10x-derived priors), Proseg, and Segger. This represents the broadest systematic comparison of segmentation algorithms in SRT to date. The evaluation is organized around three core challenges that any segmentation algorithm must navigate: producing an accurate number of cells, maximizing transcript capture within cells (minimizing false negatives), and maintaining cell purity (minimizing false positives from transcript misassignment). Transcript capture was found to be correlated with improved downstream metrics but comes at the cost of slightly reduced purity, though the magnitude of this tradeoff is tissue dependent. Overall, Proseg achieved the best balance as a generalist segmentation algorithm choice, resulting in optimal downstream metrics in most tissues, though its advantage over other methods varies by tissue.

## Results

### A multi-tissue spatial transcriptomics resource for benchmarking segmentation

To assess the impact of varying segmentation algorithms across multiple tissues, a TMA was constructed consisting of 2 mm diameter punches of formalin-fixed and paraffin-embedded (FFPE) tissue sections from 10 different mouse organs (brain, female reproductive tract [FRT], heart, kidney, large intestine, liver, lung, small intestine, spleen, and stomach). Next, SRT data were generated from it using the 10x Xenium technology platform utilizing both the multimodal segmentation kit and the Prime 5K Mouse Pan Tissue & Pathways Panel consisting of 5006 genes (**Figure 1a, Supplementary Figure 1a-c**). Segmentation using the 10x kit was performed by the onboard analyzer before being fed into pipelines for re-segmenting the data using either nuclear staining (with no extension) or one of the alternative segmentation algorithms. A total of 331,855,196 high-quality (Q-score > 20) transcripts were decoded across all tissues with 79.9% of total transcripts measured meeting this threshold. The selection of prior data was shown to be particularly important when running the Baysor algorithm ^8^, and so this method was run using either DAPI-based priors (bayNucleus) or priors derived from the 10x segmentations (bay10x).

**Figure 1.**
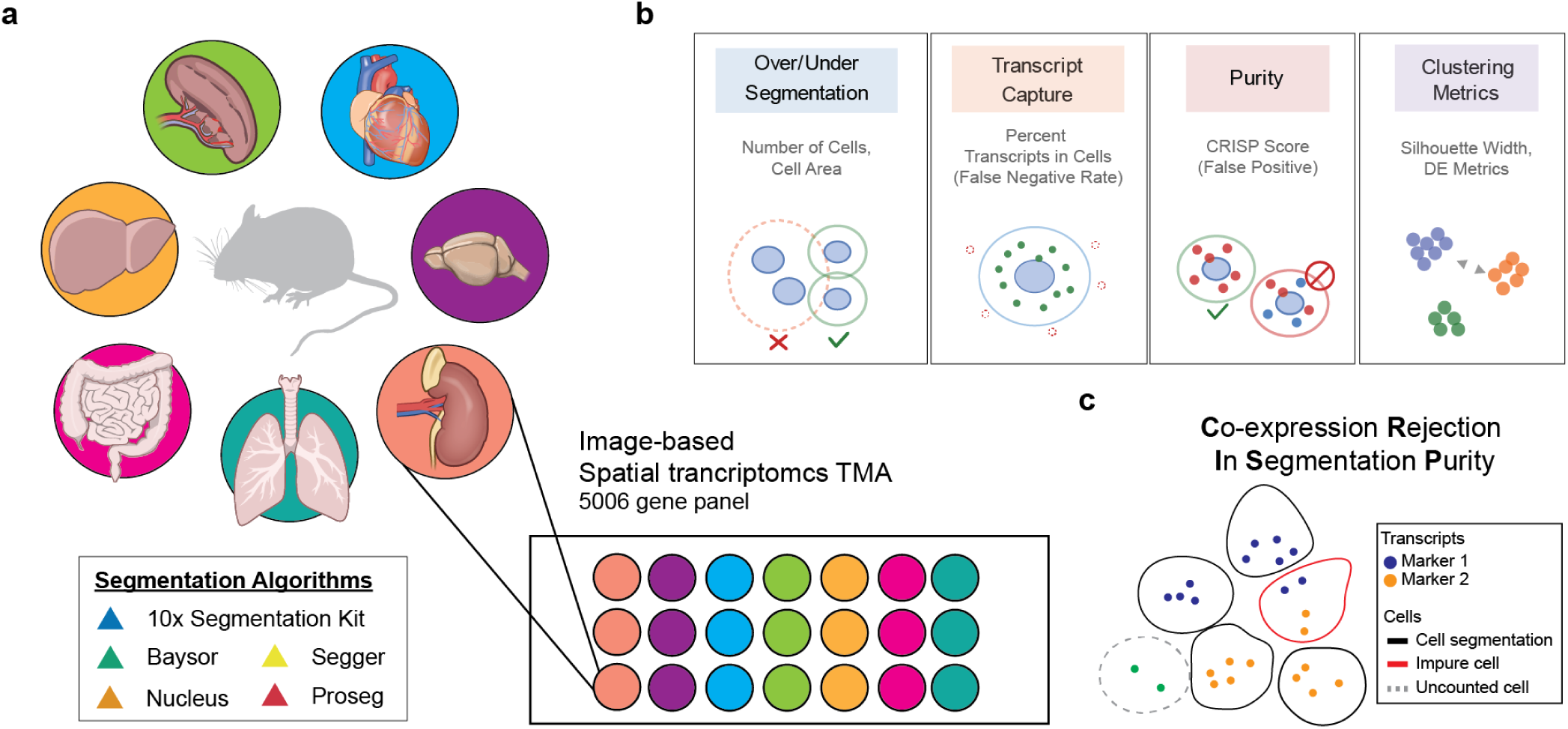
A multi-tissue spatial transcriptomics resource for benchmarking cell segmentation. **a.** Schematic of the experimental design. A tissue microarray (TMA) was constructed from formalin -fixed, paraffin-embedded (FFPE) sections of 10 mouse organs (brain, FRT, heart, kidney, large intestine, liver, lung, small intestine, spleen, and stomach) a nd profiled using the 10x Xenium platform with the Prime 5K Mouse Pan Tissue & Pathways Panel (5,006 genes). Five segmentation algorithms spanning image-based (nuclear segmentation, 10x multimodal segmentation kit) and transcript-based (Baysor with nuclear or 10x-derived priors, Proseg, and Segger) approaches were applied to the same underlying data. **b.** Overview of the four complementary metric categories used to evaluate segmentation quality: over/under-segmentation (cell counts and cell area), transcript capture (percentage of transcripts assigned to cells, reflecting false negative rates), purity (CRISP score, reflecting false positive rates from transcript misassignment), and clustering metrics (silhouette width and differential expression effect sizes). **c.** Schematic of the CRISP (Co-expression Rejection In Segmentation Purity) algorithm. CRISP quantifies segmentation purity by detecting cells that inapprop riately co-express tissue-specific mutually exclusive marker genes, which indicates chimeric cells arising from boundary errors. Cells expressing markers from two or more mutually exclusive cell types are classified as impure, while cells expressing markers from exactly one cell type are classified as pure. Cells not expressing any markers or only those not relevant to the test (green dots) are not scored.

The TMA was originally constructed as part of an independent study evaluating tissue-specific delivery of lipid nanoparticles (LNPs) encapsulating β-Galactosidase mRNA cargo in C57BL/6J mice. The array therefore includes tissue punches from both LNP-treated and vehicle control animals (**Supplementary Figure 1c–d**). Because this study focuses exclusively on the computational comparison of segmentation algorithms applied to the same underlying data, the LNP treatment status does not affect the benchmarking conclusions; all algorithms were evaluated on identical tissue sections, and any treatment-related differences in transcript profiles or cell morphology are held constant across algorithm comparisons within each tissue.

Cell segmentation errors can manifest in several ways, and no single metric is sufficient to capture overall segmentation quality. Algorithm performance was evaluated across four complementary categories of metrics (**Figure 1b**). First, over- and under-segmentation was assessed by comparing the number of cells identified by each algorithm and the distribution of cell areas, as algorithms that fragment single cells or merge neighboring cells will produce characteristic distortions in these measures. Second, transcript capture was quantified as the proportion of decoded transcripts assigned within cell boundaries, reflecting the false negative rate quantifying the extent to which transcripts fall outside segmented regions and are effectively lost from downstream analysis. Third, cell purity was measured, the tendency for segmentation algorithms to incorrectly assign transcripts to neighboring cells, representing the rate of false positive errors in the segmentation process. Lastly, integrated downstream readouts were evaluated, sensitive to systematic segmentation biases which can degrade expression profiles across the data. These metrics measure clustering quality using silhouette width scores or changes in differential expression effect sizes (log2 fold-change and percent difference between clusters) which are central attributes for many downstream analysis goals.

To accurately measure cell purity, the CRISP method was developed to quantify the rate at which segmented cells co-express mutually exclusive marker genes, an indicator of chimeric cells arising from boundary errors (**Figure 1c**). CRISP utilizes tissue-specific marker lists derived from expert knowledge or single-cell reference datasets and does not require cell type annotations or manual ground truth segmentations as in other benchmarks or tools. Marker pairs are selected such that each gene (or small gene set) is expressed at sufficient levels in the spatial data, ensuring flexibility across different SRT panel sizes and platforms. CRISP was implemented as open-source R (crispRSeg) and Python (crispySeg) packages which are available for download at (github.com/santangelo-lab/CRISP).

### CRISP detects segmentation impurity through mutually exclusive marker co-expression

CRISP requires selection of a detection threshold — the expression level at which a gene is counted as present in a cell. This parameter is critical because count values for spatially-decoded transcripts vary widely across platforms, gene panels, and natural gene expression levels. To guide threshold selection, simulation studies were conducted examining how CRISP performance depends on the interaction between threshold choice and both marker expression level and per-cell gene count. The CRISP score was calculated on a range of simulated datasets with levels of true impurity ranging from 0-30%, and sensitivity, specificity, and accuracy were computed across varying detection threshold levels. As expected, when the expression threshold was increased, the sensitivity of CRISP decreased, while specificity increased inversely (**Figure 2a**). Simulations at a mean expression level of 4 counts per cell showed that F1 score was highest when the detection threshold was set to approximately half the mean marker expression (thresholds of 2–3). When marker expression is higher (simulated at mean values of 10 or 50 counts per cell) peak accuracy occurred at higher threshold values (**Supplementary Figure 2a, 2b**). Simulations run with a mean expression of 10 had the highest accuracy when setting the threshold between 4-5, while increasing marker expression to 50 resulted in the peak accuracy at a threshold value of 10. Comparing simulated CRISP estimates of purity to true values showed that setting high detection thresholds when using lowly expressed markers can artificially inflate purity values, whereas setting low thresholds for highly expressed markers will artificially deflate purity (**Figure 2b**). All told, these results suggest that expected marker expression needs to be considered when setting the threshold parameter values as these need to be well matched to generate the most accurate estimates of purity.

**Figure 2.**
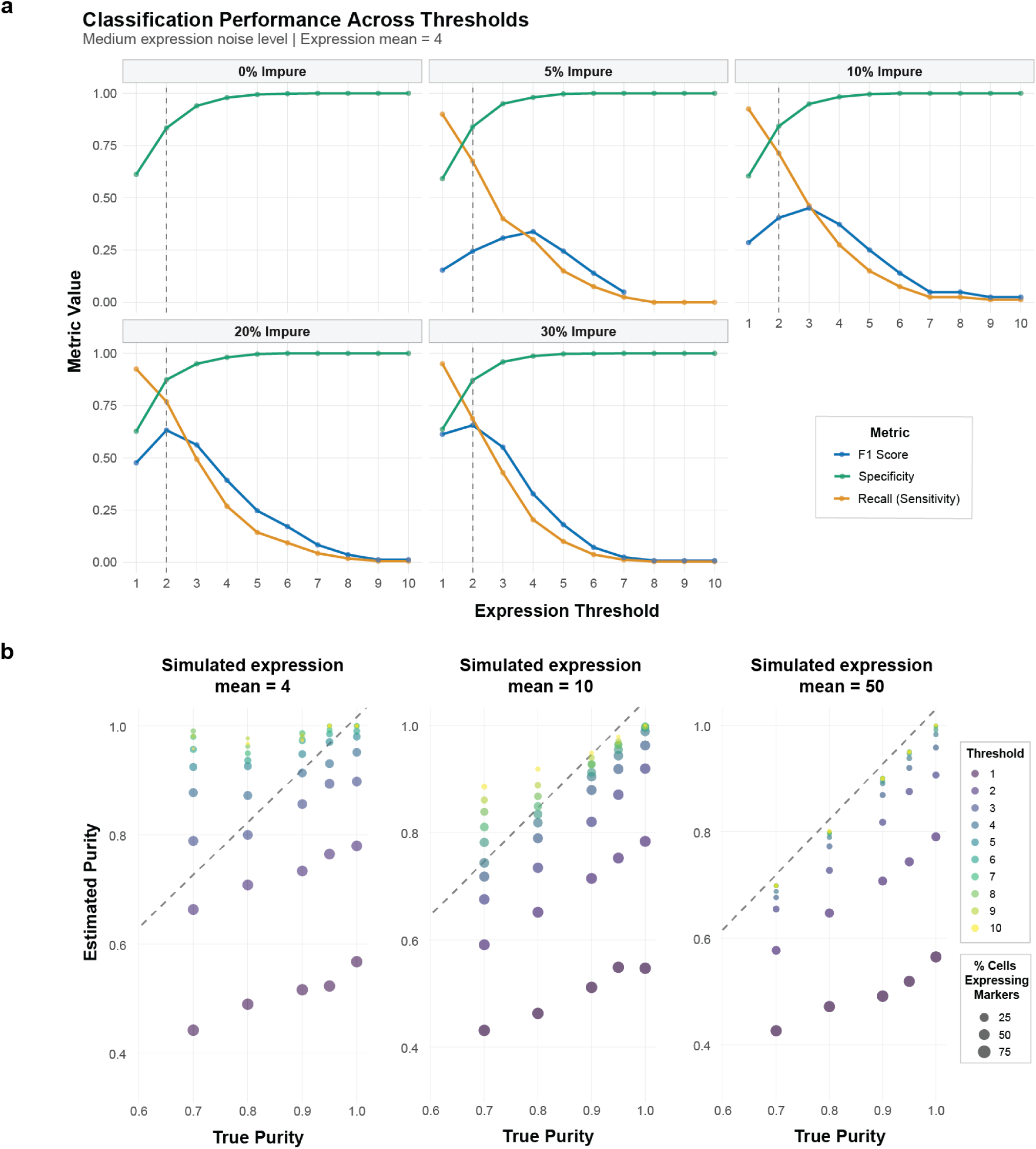
Simulation studies validate CRISP purity estimation across detection thresholds. **a.** Classification performance of CRISP across detection thresholds at varying levels of true impurity (0%, 5%, 10%, 20%, and 30%) for simulated datasets with mean marker expression of 4 counts per cell and medium expression noise. F1 score (blue), recall/sen sitivity (orange), and specificity (green) are shown for each threshold value (1 –10). **b.** Estimated versus true purity for three simulated mean marker expression levels (4, 10, and 50 counts per cell) across a range of detection thresholds (color scale). Po int size reflects the percentage of cells expressing marker genes at each threshold. The diagonal dashed line indicates perfect estimation. n = 1,0 00 cells per simulation.

Next, the sensitivity of the CRISP method required determination in relation to variations in sample size. Data were isolated from four tissue types (heart, liver, small intestine, and spleen) found within the TMA, and the number of cells expressing the marker panel genes was randomly downsampled to varying degrees. CRISP purity scores were computed at each sample size across 20 repeated subsamples to assess both the accuracy and precision of the estimates (**Figure 3a**). Purity scores were tissue-specific, with liver and spleen exhibiting higher purity (∼0.98–0.99) than heart (∼0.96–0.97) and small intestine (∼0.94–0.95), consistent with differences in tissue morphology and cell packing density. Notably, the median purity estimate remained stable across sample sizes, with scores at the smallest sample sizes closely approximating those obtained from the full dataset in each tissue. Variance in the estimates narrowed progressively with increasing sample size, as reflected by decreasing interquartile ranges across successive downsampling levels, and was negligible beyond approximately 2,500–5,000 cells, suggesting that CRISP scores are robust to the range of dataset sizes commonly encountered in spatial transcriptomics experiments. These results indicate that CRISP provides reliable purity estimates even in moderately sized datasets and that meaningful comparisons can be made across tissues with different numbers of marker-expressing cells.

**Figure 3.**
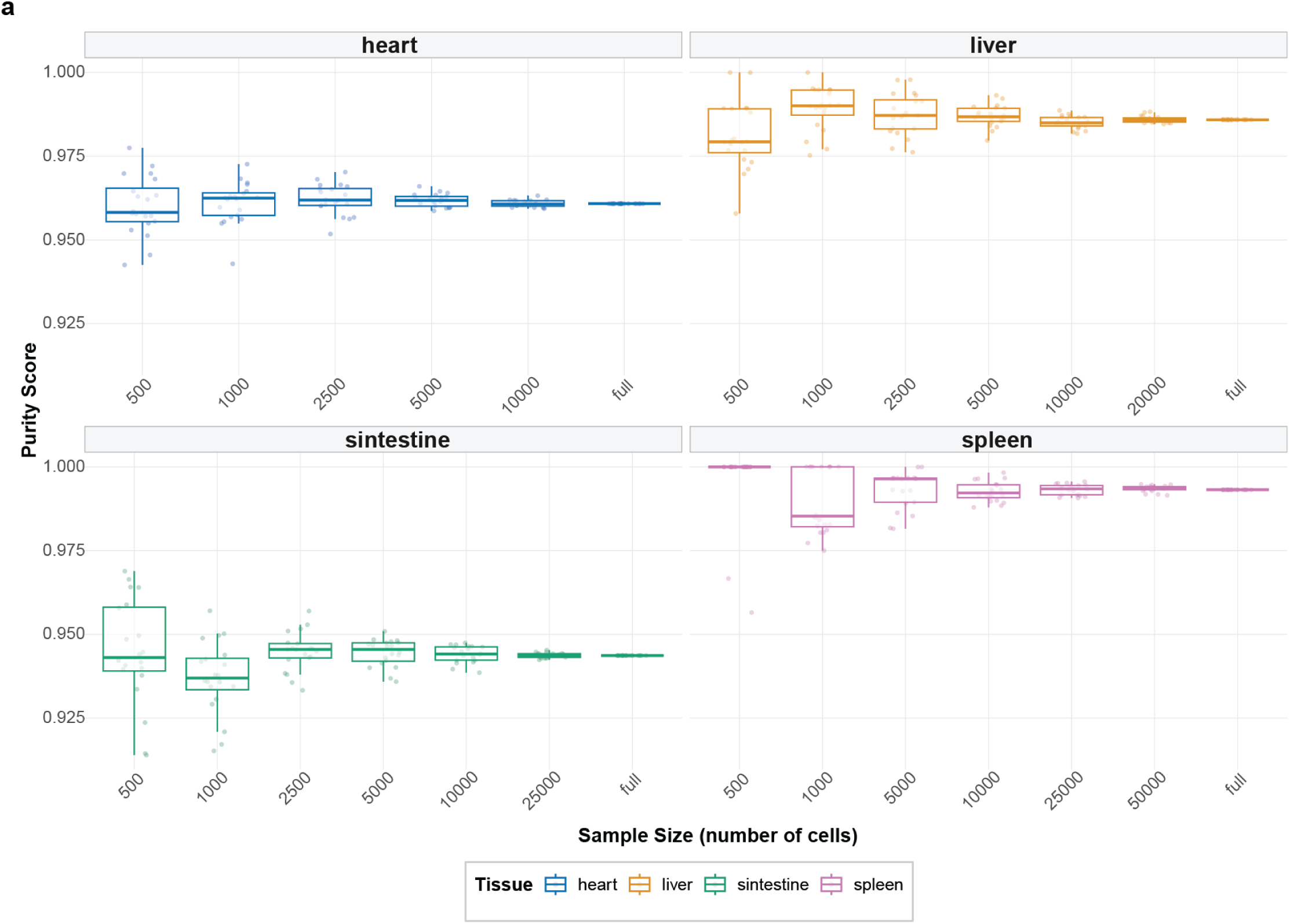
CRISP purity estimates are stable across sample sizes. **a.** Box plots of CRISP purity scores across downsampled datasets at varying sample sizes for four tissues: heart, liver, small in testine (sintestine), and spleen. For each tissue, cells were randomly downsampled to the indicated sample sizes, and CRISP was computed across 20 independent subsamples per size. Box plots show the median (center line), first and third quartiles (box limits), a nd 1.5× interquartile range (whiskers). Detection threshold = 3 counts. Data are from the 10x multimodal segmentation kit output.

### Cross-tissue benchmarking reveals a transcript capture–purity tradeoff

In the absence of manually curated ground truth segmentations, DAPI-derived nuclear boundaries were used as the most conservative and morphologically grounded reference for expected cell counts. The number of cells identified by each algorithm varied dramatically, with the two Baysor configurations producing the largest departures: both bay10x and bayNucleus consistently generated 2–7x more cells than nuclear segmentation across all tissues (**Figure 4a**). Using the 10x segmentations as prior data in the Baysor algorithm slightly reduced the number of cells segmented; however, both Baysor configurations yielded substantial oversegmentation. In contrast, Proseg, Segger, and the 10x segmentation kit produced cell counts within approximately 2-fold of nuclear estimates in most tissues, with Segger producing slightly fewer segmented cells across most tissues. The degree of Baysor oversegmentation was tissue-dependent, with the most extreme inflation observed in liver and heart, and the least in spleen tissue.

**Figure 4.**
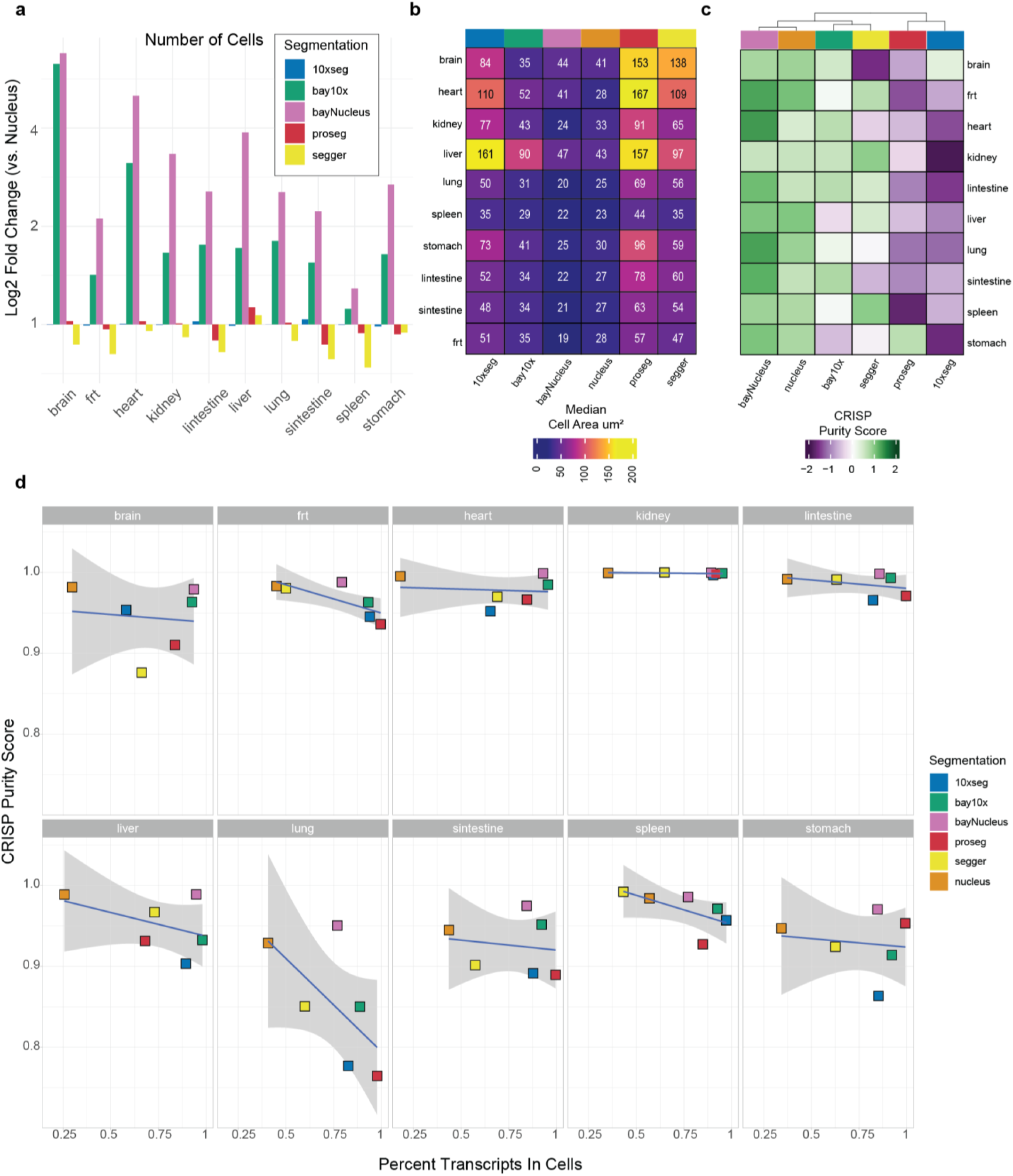
Cross-tissue benchmarking reveals a tissue-dependent transcript capture–purity tradeoff. **a.** Cell counts for each algorithm expressed as log2 fold-change relative to the nuclear segmentation baseline within each tissue. **b.** Heatmap of median cell area (µm²) across all tissue–algorithm combinations. Values are displayed within each cell. **c.** Heatmap of CRISP purity scores across all tissue–algorithm combinations, z-score scaled within each tissue. Color scale represents z-score deviations from the tissue mean. **d.** Scatter plots of CRISP purity score versus percent of transcripts assigned to cells for each tissue, with each point representing one algorithm. Trend lines represent linear fits estimated using a generalized linear model. Sha ded regions indicate 95% confidence intervals. Algorithms are color-coded; n = 6 algorithms per tissue.

Algorithms also differed substantially in the size of the cells they produced (**Figure 4b**). Nuclear segmentation yielded the smallest median cell areas across all tissues, as expected given the absence of cytoplasmic extension. BayNucleus cells were similarly compact. Proseg and Segger generated the largest cell areas, reflecting more aggressive strategies to extend boundaries beyond the nucleus to capture cytoplasmic transcripts.

Notably, the relative ranking of algorithms by cell area was largely conserved across tissues, though absolute areas varied considerably with liver cells among the largest regardless of algorithm, while spleen and lung cells were consistently the smallest, reflecting genuine differences in cell morphology.

These differences in median cell area were strongly correlated with median transcripts per cell across all tissue–algorithm combinations (R = 0.87, p < 2.2 × 10⁻¹⁶; **Supplementary Figure 3a**), likely due to the fact that algorithms which draw larger boundaries capture proportionally more transcripts. This relationship has important implications for downstream analysis: median transcripts per cell was positively correlated with clustering quality metrics including mean silhouette width and differential expression effect sizes (mean percent difference between clusters), while being negatively correlated with CRISP purity scores (**Supplementary Figure 3b**). Importantly, these relationships were not uniform across tissues (**Supplementary Figure 3c**): the negative correlation between transcripts per cell and CRISP purity was strongest in the lung, large intestine, and FRT, suggesting that the severity of this capture–purity tradeoff is modulated by tissue architecture.

Next, how these tradeoffs manifested in CRISP purity scores was examined across all tissue–algorithm combinations (**Figure 4c**). Nuclear segmentation and bayNucleus achieved the highest purity scores across most tissues, consistent with their conservative, compact boundaries. Proseg and the 10x segmentation kit exhibited the lowest purity in most tissues, reflecting the cost of their more expansive segmentation strategies. However, the magnitude of purity differences between algorithms varied substantially by tissue: brain, lung, and liver showed the widest separation between the highest- and lowest-purity algorithms, while spleen and small intestine showed more compressed ranges, suggesting that some tissue architectures are inherently more tolerant of boundary expansion.

The capture–purity tradeoff was directly visualized by plotting CRISP purity against the percent of transcripts assigned to cells for each tissue (**Figure 4d**). In most tissues, a negative trend was apparent, with algorithms achieving higher transcript capture rates tending to have lower purity scores. However, the slope and tightness of this relationship varied by tissue. Brain, lung, FRT, and liver showed steep tradeoffs, where gains in transcript capture came at substantial purity cost, while kidney, heart, intestine (both large and small), and spleen showed relatively flat relationships where algorithms could increase capture with more modest purity penalties. Nuclear segmentation consistently occupied the high-purity, low-capture corner of these plots, while Proseg and the 10x segmentation kit tended toward higher capture with lower purity (**Supplementary Figure 4a**). Notably, Segger frequently achieved intermediate positions in both metrics, suggesting a moderate balance between capture and purity across tissues. Radar plots summarizing all four metric categories across tissues confirmed that no single algorithm dominated across all dimensions (**Supplementary Figure 4b**), though Proseg consistently achieved the strongest combined performance in transcript capture, clustering quality, and differential expression metrics across the majority of tissues, despite its lower purity scores. One notable exception was liver tissue, where Proseg struggled to capture transcripts and failed to exceed the other methods in any other metric.

### Segmentation choice impacts clustering quality and differential expression

To examine how the tradeoffs identified above manifest in a specific downstream analysis context, a detailed case study was performed in mouse brain tissue, one of the most highly-studied tissue types in the SRT literature and a tissue type that has been used in the development of many different segmentation algorithms. Unsupervised clustering and UMAP visualization revealed differences in the structure and resolution of cell populations across algorithms (**Figure 5a**). The number of clusters identified using a constant clustering resolution value of 0.2 ranged from 9 (Segger) to 21 (Proseg), with nuclear segmentation producing 11 clusters, bayNucleus producing 15 clusters, bay10x producing 17 clusters, and the 10x segmentation kit producing 15.

**Figure 5.**
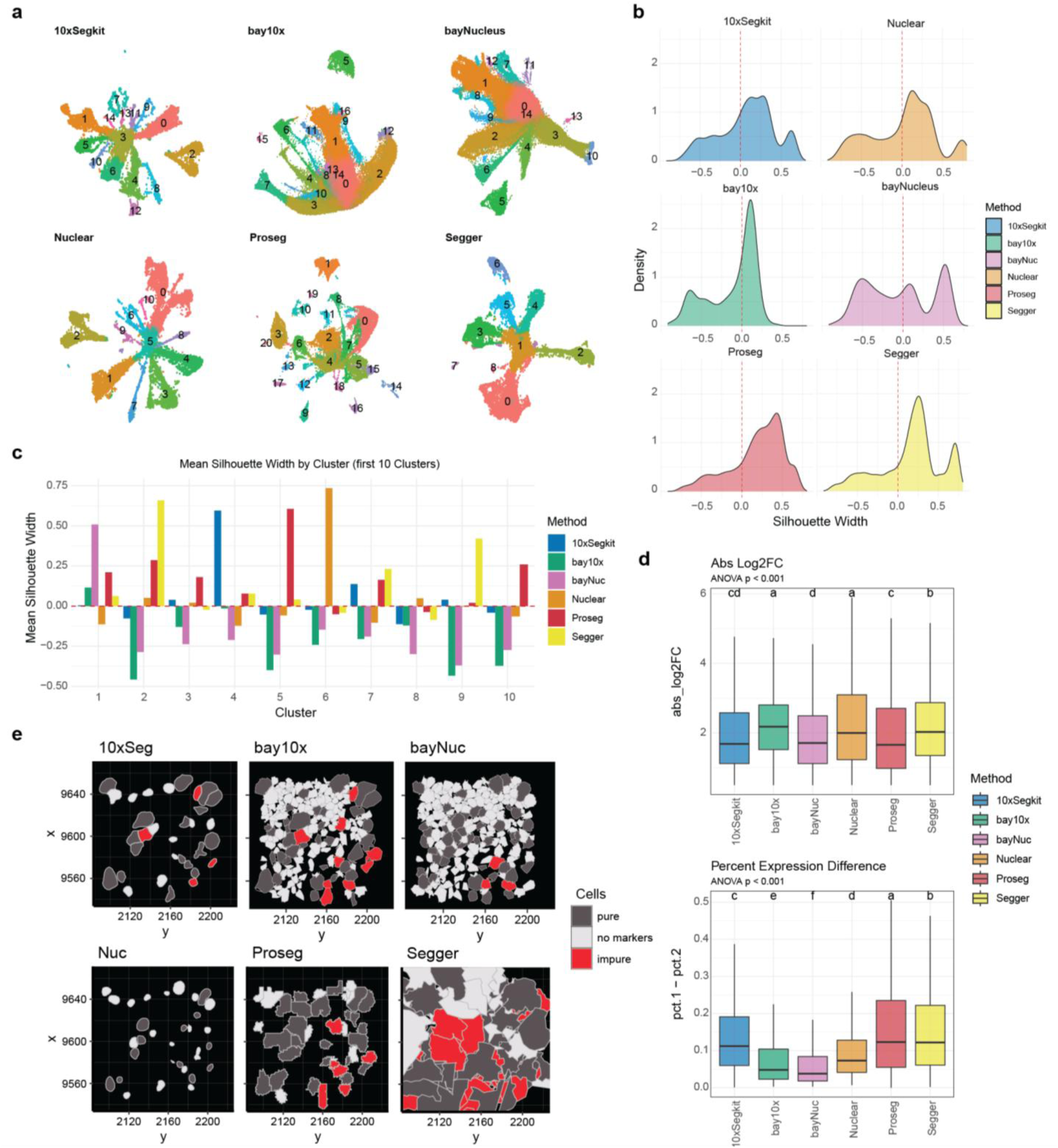
Segmentation algorithm choice shapes clustering resolution and differential expression in mouse brain. **a.** UMAP embeddings of mouse brain tissue for each segmentation algorithm, colored by unsupervised Louvain cluster identity (resolution = 0.2). The number of clusters ranges from 9 (Segger) to 21 (Proseg). **b.** Silhouette width distributions across all cells for each algorithm. Euclidean distances were computed in PCA space on a subsample of up to 5,000 cells. **c.** Mean silhouette width by cluster for the first 10 clusters across all algorithms. **d.** Differential expression metrics for cluster marker genes. Left: distributions of absolute log2 fold-changes across algorithms (ANOVA p < 0.001); compact letter displays indicate distinct pairwise significance groups (Tukey’s HSD, p < 0.05). Right: distributions of percent expressed difference (pct.1 − pct.2) between cl usters of interest and all other clusters (ANOVA p < 0.001). Marker genes were identified using Seurat FindAllMarkers (log2FC threshold = 0.5, Wilcoxon rank -sum test, Bonferroni-adjusted p < 0.05). **e.** Spatial visualization of CRISP purity classifications in a representative brain region colored by purity status: pure (single marker type detected), impure (co -expression of mutually exclusive markers), or no markers (below detection threshold).

The quality of clustering was further assessed by examining silhouette width distributions for each algorithm across all cells (**Figure 5b**). Proseg and Segger produced distributions that shifted toward positive values relative to the other methods, indicating tighter within-cluster coherence and greater separation between clusters. Nuclear segmentation and the 10x segmentation kit showed broader distributions centered closer to zero, with a substantial proportion of cells exhibiting negative silhouette widths, a signature of ambiguous cluster assignments likely arising from noisier expression profiles due to limited transcript capture^15^. Per-cluster analysis of mean silhouette width confirmed that Proseg achieved the highest or near-highest scores across most clusters, while Baysor segmentation consistently performed poorly regardless of prior data used (**Figure 5c**).

Differential expression analysis reinforced these findings (**Figure 5d**). The distribution of absolute log2 fold-changes for cluster marker genes differed significantly across algorithms (ANOVA p < 0.001), with bay10x, Nuclear, and Segger producing the largest effect sizes. A similar pattern was observed for the percent expressed difference between clusters, where Proseg and Segger yielded the most distinctive marker expression profiles (**Figure 5d**). These results indicate that log2FC may be most sensitive to high purity approaches like nuclear whereas algorithms capturing more transcripts per cell generate expression profiles with greater discriminative power, enabling cleaner identification of cell type–specific markers — a practical advantage for atlas-building and cell type annotation efforts. However, these gains in clustering and differential expression quality come at a measurable cost to segmentation purity. Spatial visualization of CRISP classifications in a representative brain region revealed the nature of this tradeoff at the cell-level resolution (**Figure 5e**). Nuclear segmentation produced the fewest impure cells, with most boundaries tightly enclosing individual nuclei and leaving substantial interstitial space unassigned. The 10x segmentation kit expanded boundaries modestly, introducing a small number of impure cells at boundaries between cell types. Impurities in the Baysor algorithms largely followed the prior dataset used, however each produce large quantities of small cells not expressing sufficient quantities of measured markers. Proseg, while achieving the strongest clustering metrics, showed a markedly higher density of impure cells, particularly in regions where morphologically distinct cell types were closely interspersed. Segger exhibited an intermediate purity profile, while drawing the most large, irregular cell boundaries – possibly due to the fact that, like Baysor, the algorithm relies primarily on transcript assignment instead of boundary drawing. These qualitative patterns hold true among all assayed tissues (**Supplementary Figure 5**). Overall, these spatial patterns illustrate that the algorithms which most effectively improve downstream expression analysis do so in part by drawing the most expansive *and* plausible boundaries, but they are more prone to incorporating transcripts from neighboring cells — particularly in the densely packed, heterogeneous cytoarchitecture of the brain.

### Integrated performance assessment and tissue-specific recommendations

All of benchmarking metrics were compiled into a unified visualization for each tissue, with values scaled across algorithms within each metric category (**Figure 6**). This integrated assessment reinforced a central finding of our benchmark: no single algorithm dominated across all metrics in any tissue. Instead, each method exhibited a characteristic performance profile reflecting its underlying design strategy, and the relative advantages of each approach varied with tissue context. Proseg, however, achieved the highest average rank across tissues in the majority of individual metrics, excelling in transcript capture, clustering quality, and differential expression effect sizes simultaneously (**Figure 7**). Its consistent performance across these categories, which together determine the quality of most common downstream analyses including cell type identification, marker gene discovery, and expression quantification, positions it as the strongest average performer in our benchmark.

**Figure 6.**
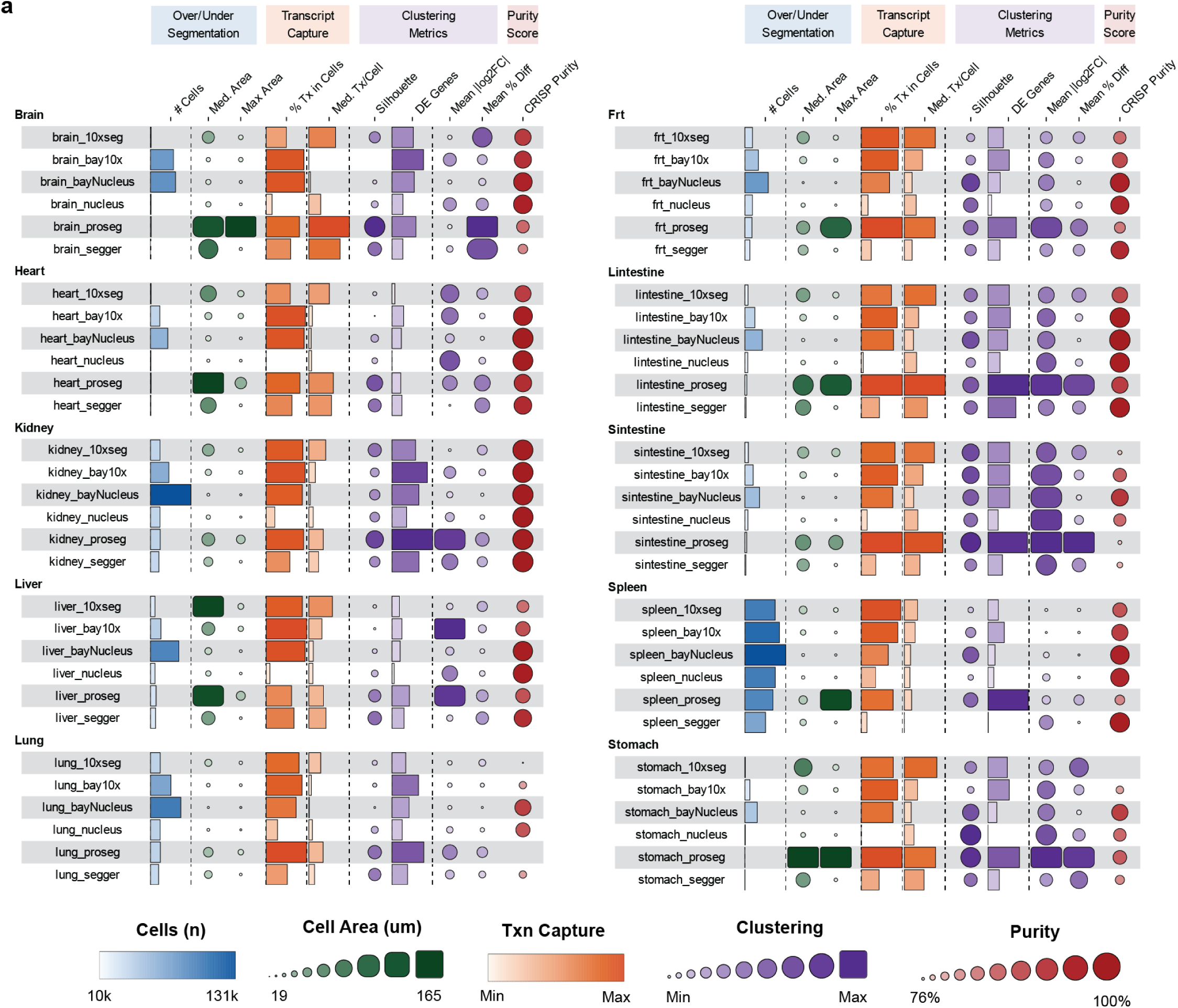
Integrated performance profiles reveal tissue-specific algorithm strengths and tradeoffs. **a.** Comprehensive heatmaps of benchmarking metrics across all algorithm–tissue combinations for ten tissues, organized in two panels (brain, heart, kidney, liver, lung in the first panel; FRT, large intestine, small intestine, spleen, stomach in the second). Metrics are grouped into five categories: over/under-segmentation (number of cells, median cell area, maximum cell area), transcript capture (percent transcripts in cells, median transcripts per cell), clustering metrics (mean silhouette width), differen tial expression metrics (number of DE genes, mean absolute log2 fold-change, mean percent difference), and purity (CRISP purity score). Values are min–max scaled within each metric and tissue to enable cross-algorithm comparison. Color scale ranges from low (blue) to high (red) relative to performance within each tissue. Cell counts range from approximately 10,000 to 131,000; cell areas from 19 to 165 µm²; transcript capture from 76% to 100%.

**Figure 7.**
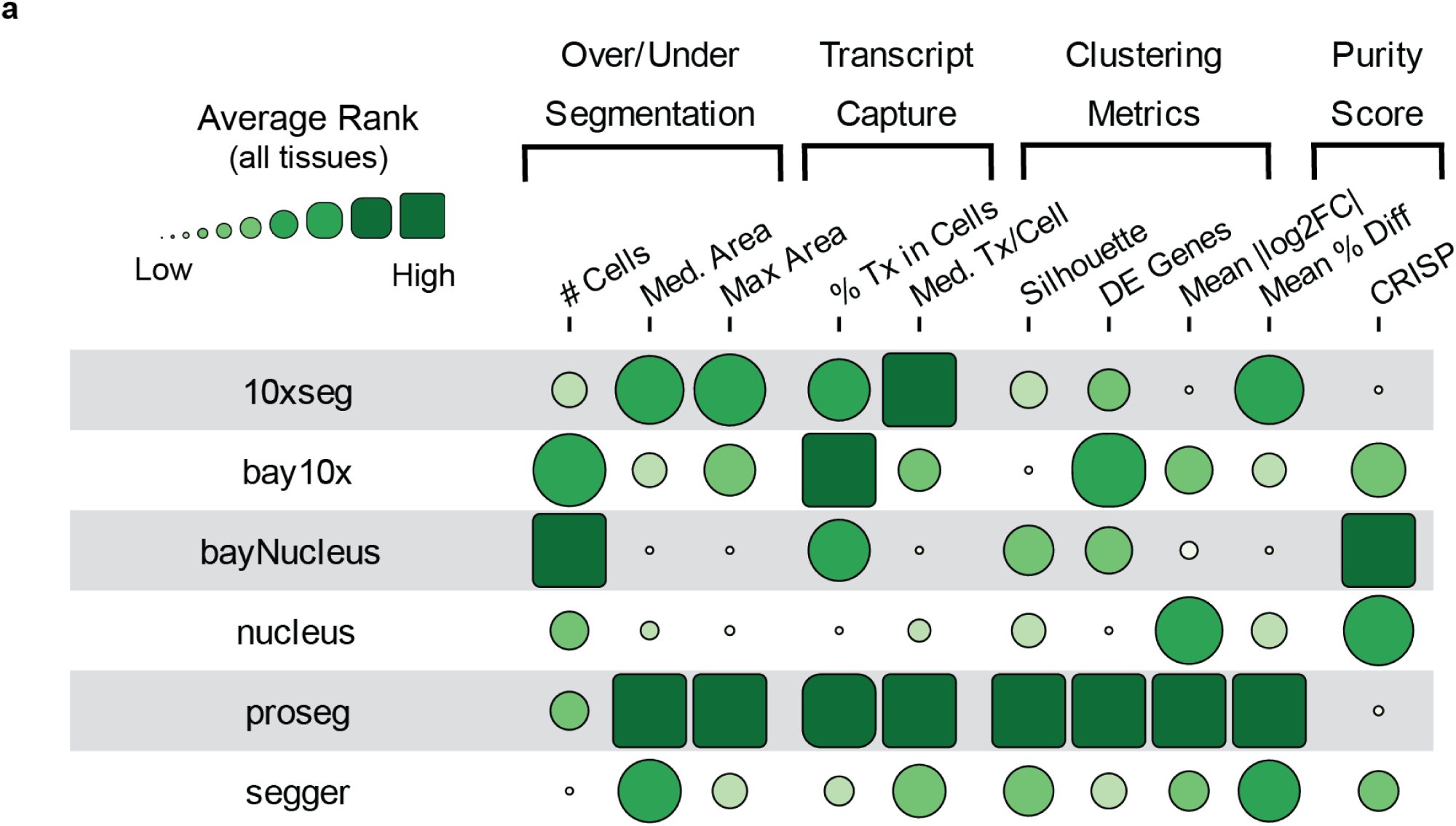
Proseg achieves the strongest average performance across tissues. **a.** Heatmap of average metric ranks across all ten tissues for each algorithm. For each tissue, algorithms were ranked within each metric (1 = best, 6 = worst), and ranks were averaged across tissues. Metrics are organized into the same five categories as Figure 6: over/under-segmentation, transcript capture, clustering, differential expression, and purity. Color scale ranges from low (light) to high (dark) average rank.

In liver, Proseg and Segger showed notably lower transcript capture compared to the 10x segmentation kit despite producing comparable cell areas, suggesting that the unique sinusoidal architecture and large hepatocyte cell bodies of liver tissue may favor the image-based boundary detection strategy employed by the 10x kit over the transcript-density-based approaches used by Proseg and Segger. Spleen also presented a notable exception, where the compressed range of performance across all algorithms and metrics suggests that the densely packed, relatively homogeneous, lymphoid architecture of this tissue is tolerant of segmentation approach, diminishing the practical advantage of any single method. These tissue-specific departures underscore that while Proseg serves as the strongest general-purpose recommendation, researchers working with specific tissue types should consult the per-tissue performance profiles to confirm that the segmentation algorithm choice is appropriate for their context.

## Discussion

Here, two principal contributions are presented to the spatial transcriptomics community. First, the most comprehensive independent benchmark of cell segmentation algorithms for image-based spatial transcriptomics to date is provided, comparing five commonly used segmentation strategies across ten mouse tissues profiled with a 5,006-gene Xenium panel. Second, CRISP is introduced, a fully packaged software tool, available in both R and Python, for quantifying segmentation purity in image-based spatial transcriptomics data using biologically informed, tissue-specific mutually exclusive markers. Together, these contributions address a critical gap in the field: despite the importance of segmentation quality for downstream analysis, no prior study has independently evaluated segmentation algorithms at this scale or provided accessible, packaged tools for researchers to assess purity in their own data.

The evaluation was organized around three critical attributes of cell segmentation quality: accuracy of cell number estimation (over- or under-segmentation), minimization of false negatives (transcript capture), and minimization of false positives (cell purity). These metric categories are not independent. Algorithms that draw more expansive cell boundaries will capture more transcripts per cell but risk incorporating molecules from neighboring cells, creating a fundamental tradeoff between transcript capture and cell purity whose severity depends on tissue morphology. Transcript capture was found to be negatively correlated with CRISP purity scores, but at the same time positively associated with multiple important downstream analysis metrics including the ability to resolve distinct cell clusters and the detection of differentially expressed genes. Downstream metrics such as clustering silhouette widths and differential gene expression effect sizes thus serve as integrated readouts of the balance between false negative and false positive rates, rather than independent evaluation axes. These correlations reveal a fundamental tension in segmentation: expanding cell boundaries improves transcript recovery and strengthens the expression signatures that drive clustering and marker gene detection while simultaneously increasing the risk of incorporating transcripts from adjacent cells.

Several algorithm-specific patterns emerge clearly from the multi-tissue benchmark. Baysor consistently oversegments, producing two- to seven-fold more cells than nuclear segmentation across tissues regardless of prior data input, a finding consistent with observations from the original Proseg benchmarks^9^. This oversegmentation appears to reflect cell fragmentation rather than appropriate boundary extension since Baysor variants perform particularly poorly on integrated downstream metrics despite achieving favorable purity and capture scores; the fragmentation of gene signatures within individual cells likely undermines the coherent expression profiles needed for robust clustering and differential expression analysis. Conversely, Proseg consistently achieves a favorable balance between transcript capture and purity across most tissues. The 10x default segmentation kit, despite leveraging multimodal immunofluorescence staining, achieves competitive transcript capture rates but consistently underperforms transcript-based methods in clustering quality and cell purity across most tissues. It is worth noting that the performance of the 10x segmentation kit is inherently dependent on the quality of the underlying immunofluorescence stains, which could vary with tissue fixation conditions, staining protocol optimization, and tissue-specific differences. The relatively poor performance observed here may therefore reflect the challenges of achieving uniformly high-quality staining across ten diverse tissues using a single standardized protocol, suggesting that image-based segmentation approaches may require tissue-specific staining optimization to compete with transcript-based methods.

A central finding of this study is that the severity of the transcript capture–purity tradeoff is highly tissue-dependent. Some tissues, such as kidney, exhibit a relatively modest exchange rate between increased transcript capture and purity loss, whereas others, particularly lung, show much steeper purity costs for equivalent gains in capture. This tissue dependence likely reflects the underlying cellular architecture of each organ. Tissues composed of tightly packed, small cells from many distinct lineages, such as in lung tissue, present a far more challenging environment for segmentation than tissues with larger, well-separated cells of fewer types, where boundary errors are less likely to merge transcriptionally distinct neighbors. The initial hypothesis was that different algorithms would perform best in different tissues, but the results paint a more nuanced picture. While the magnitude of performance differences varies substantially by tissue, the relative ranking of algorithms is largely consistent, with Proseg achieving the highest average ranks across tissues in transcript capture, over/under segmentation accuracy, and downstream clustering and differential expression metrics. This pattern suggests that its transcript-density-based boundary estimation effectively reduces false negatives without proportionally increasing false positives — a tradeoff that most tissues tolerate well. However, it is notable that in several tissues, including spleen, stomach, and small intestine, performance differences between the top algorithms were compressed, indicating that the practical benefit of algorithm choice is context-dependent. Nonetheless, this suggests that the underlying principles governing transcript-based segmentation may be more broadly applicable for achieving high quality segmentations across multiple tissues compared to image/staining-based methods.

Several approaches for measuring segmentation purity in spatial transcriptomics have been proposed in recent work, and it is important to understand how CRISP relates to and differs from these alternatives. The MECR (mutually exclusive coexpression rate) introduced by Hartman and Satija^16^ is the most direct conceptual antecedent of CRISP, as both quantify inappropriate co-expression of markers that should be confined to distinct cell types. However, the two methods differ in a few important respects. MECR computes a per-gene-pair co-expression rate similar to CRISP but averages this rate across all mutually exclusive pairs identified from a single scRNA-seq reference using automated differential expression filtering. CRISP instead operates at the cell level, classifying each individual cell as pure or impure based on whether it co-expresses pre-defined markers from more than one cell type, and computing purity as a proportion of marker-positive cells. This cell-level approach enables spatial mapping of impure cells and identification of which specific cell-type interfaces are most affected by segmentation errors. Additionally, whereas MECR was applied with a single universal marker set in mouse brain tissue, CRISP uses tissue-specific curated marker panels, supports compound markers that sum multiple genes to increase sensitivity for cell types with low individual marker expression, and incorporates threshold sensitivity analyses with automated diagnostic warnings to guide appropriate parameter selection. CRISP is also implemented and released as open-source software in both R and Python. While the underlying biological principle is shared, these differences make CRISP a distinct tool designed for cross-tissue segmentation benchmarking rather than cross-platform specificity assessment.

CellSPA, another segmentation assessment tool introduced alongside BIDCell^17^, takes a broader benchmarking approach that includes a positive marker purity metric. CellSPA calculates precision, recall, and F1 scores for both positive and negative cell type markers, combining them into a composite purity F1 score. While comprehensive, this approach has two important requirements that limit its accessibility: it requires prior cell type labels for all segmented cells, and it depends on a matched single-cell RNA-seq reference dataset to define positive and negative marker sets. In contrast, CRISP operates directly on expression data without requiring cell type annotations, making it applicable earlier in the analysis workflow before cell typing has been performed and avoiding the circularity of evaluating segmentation quality using labels that are themselves affected by segmentation errors. Similarly, the NMP (negative marker purity) metric introduced by Salas et al.^7^ quantifies the excess presence of negative markers across cell types relative to a single-cell reference, but like CellSPA, requires both cell type annotations and a reference dataset, and was presented as a metric within a broader analysis pipeline rather than as a standalone tool. Plummer et al.^14^ assessed segmentation purity through robust cell type decomposition (RCTD), borrowed from lower-resolution sequencing-based SRT methods where they treat each cell as a mixture of pure cell types using matched snPATHO-seq data as a reference, an approach that provides a rich view of transcript mixing but requires paired high-quality reference data that is not available for most experiments. Among these approaches, CRISP is distinguished by its combination of label-free operation, tissue-specific marker infrastructure, and availability as packaged software, which together lower the barrier to routine purity assessment in the absence of ground truth annotations or paired reference data.

This study has several limitations. The benchmark is restricted to a single platform (10x Xenium), and the extent to which these findings generalize to other spatial transcriptomics platforms such as Vizgen MERSCOPE and NanoString CosMx remains to be determined, though the Proseg developers have demonstrated strong cross-platform performance in their own evaluations^9^. CRISP requires a priori knowledge of mutually exclusive markers for each tissue of interest, and the quality of purity estimates is inherently dependent on appropriate marker selection. By characterizing the effect of marker choice and detection threshold through simulation studies as well as incorporating these findings into automated warnings in the CRISP package, the aim was to enable users to select the best possible markers for their tissue; however, ultimately this choice is highly dependent on individual studies. The focus was on the most commonly used and accessible methods that represent the broad categories of approaches currently used in cell segmentation in an effort to maximize practical relevance for the typical Xenium user, but future benchmarking efforts should expand the algorithmic coverage.

Several future directions emerge from this work. Extending the CRISP framework to human tissue datasets and other spatial transcriptomics platforms would broaden its utility and enable cross-platform segmentation benchmarking. Integration with automated marker selection pipelines could further reduce the barrier to entry for researchers unfamiliar with tissue-specific marker biology. The tissue-dependent tradeoff patterns identified could potentially be leveraged to develop adaptive segmentation strategies that adjust boundary expansion based on local tissue architecture. Furthermore, the complementary strengths of CRISP (diagnostic evaluation) and approaches like cellAdmix^4^ (post hoc correction) suggest that combining these tools in an integrated workflow could yield higher-quality spatial transcriptomics datasets than either approach alone. In conclusion, the results demonstrate that segmentation algorithm choice has substantial, tissue-dependent effects on spatial transcriptomics data quality, and that these effects propagate through all stages of downstream analysis. Rather than identifying a universally optimal algorithm, a framework was provided for making informed, context-dependent decisions. CRISP provides the quantitative tools needed to navigate these tradeoffs, and the comprehensive benchmark establishes the empirical foundation for tissue-specific algorithm recommendations. As spatial transcriptomics data continue to grow in scale and complexity, principled evaluation of segmentation quality will be essential for ensuring that biological conclusions are robust to the technical choices made during data processing.

## Methods

### Tissue preparation and TMA dataset generation

The primary purpose of the TMA array used in this study was to assess tissue-specific delivery of lipid nanoparticles delivering β-Galactosidase mRNA cargo. Tissue sections selected for the array were made up of replicates including LNP-treated and control animals.

### mRNA synthesis, LNP formulation, and administration

β-Galactosidase mRNA was synthesized by in vitro transcription using N¹-methyl-pseudouridine modifications as previously described^18,19^. Lipid nanoparticles (LNPs) were formulated with cKK-E12 ionizable lipid at a 20:1 lipid-to-mRNA mass ratio using microfluidic mixing and characterized by dynamic light scattering and RiboGreen encapsulation assay. All animal procedures were approved by the Emory University IACUC (protocol PROTO202400000121). C57BL/6J mice (8 weeks old; Jackson Laboratory) received a single retro-orbital injection of LNPs encapsulating β-Galactosidase mRNA (40 µg, 2 mg/kg) or vehicle (DPBS) control. Tissues were harvested 24 hours post-injection following cardiac perfusion with formalin fixative.

### Tissue collection

Animals were sedated with isoflurane and then perfused by cardiac puncture with 10ml of 1x DPBS followed by 10ml of 10% neutral buffered formalin. Tissues were harvested and stored in 10% neutral buffered formalin for 24 hours and then paraffin embedded by the Emory Winship Cancer Tissue and Pathology core. Following tissue harvesting, sections of brain, FRT (ovary), heart, kidney, large intestine, liver, inguinal lymph node, lung, small intestine, spleen, and stomach were collected and routinely processed for hematoxylin and eosin (H&E) staining. H&E slides were scanned by the Emory Winship Cancer Tissue and Pathology core on a Perkin Elmer Vectra Polaris slide scanner.

### Area Selections for TMA Construction

Immunofluorescent whole tissue scans of β-Galactosidase-stained sections were used to select circular (2mm in diameter) regions to sample for the TMA. The area with the strongest positive β-Galactosidase signal, when present, for each β-Galactosidase mRNA treated tissue was chosen. For vehicle treated tissues, the equivalent structural area to its mRNA treated counterpart was chosen to ensure comparable cell types. These selections were mapped onto the corresponding paraffin blocks of each scan. 2mm diameter cores of the selected areas were punched from the block of each sample, and re-embedded into a single block to create the array. The layout of the array was designed with 8 rows and 3 columns, with 1mm between each row and 0.5mm between each column. This provided maximum tissue area while ensuring the array would fit within the fiducial border of the Xenium slide.

### Xenium *In Situ* Hybridization

Following construction of the TMA block, sections were cut and mounted onto Xenium slides (10× Genomics) and processed according to the manufacturer’s protocol for Xenium Prime *In Situ* Gene Expression with optional Cell Segmentation Staining.

### Cell segmentation algorithms

#### Nuclear segmentation

Nuclear segmentation was performed using XeniumRanger (v3.1.1.0, 10x Genomics) resegment with an expansion distance of 0 µm, representing DAPI-derived nuclear boundaries with no cytoplasmic extension. Both boundary and interior immunofluorescence stains were disabled (--boundary-stain=disable, --interior-stain=disable), ensuring segmentation relied solely on the DAPI nuclear channel. This configuration served as the most conservative baseline. Nuclear segmentation was performed on the full TMA using XeniumRanger with 32 CPU cores and 128 GB memory. The initial on-instrument nuclear detection was performed by Xenium Onboard Analysis (XOA) v3.3.0.1 (10x Genomics).

### 10x Segmentation Kit

Cell segmentation using the 10x multimodal segmentation kit was performed on-instrument by the Xenium analyzer using Xenium Onboard Analysis (XOA) v3.3.0.1 (10x Genomics). This approach combines DAPI nuclear staining with additional immunofluorescence channels targeting interior protein, cell surface, and cytoskeletal markers provided by the Xenium Prime Cell Segmentation Staining reagents (10x Genomics). See Supplementary Figure 1e for staining across TMA dataset. A machine learning model integrated into XOA uses these multimodal staining channels to delineate cell boundaries beyond the nucleus. No additional post-processing or re-segmentation was applied to the 10x segmentation output.

#### Baysor

Transcript-based cell segmentation was performed using Baysor (v0.7.1) in two configurations differing in prior segmentation data. In the nuclear prior configuration (bayNucleus), DAPI-derived nuclear segmentation masks from the XeniumRanger resegment output were provided as priors, with a minimum molecules per cell threshold of 10. In the 10x prior configuration (bay10x), cell assignments from the 10x multimodal segmentation were used as priors, with a minimum molecules per cell threshold of 30. Both configurations used a prior segmentation confidence of 0.8, polygon output in GeometryCollectionLegacy format, and included x, y, and z coordinate information for three-dimensional transcript positioning. The scale parameter (expected cell radius) was estimated automatically by Baysor from the data at runtime. To manage computational requirements, the TMA was divided into a grid of tiles and each tile was processed independently in parallel. Tiles containing fewer than 100 transcripts were excluded from segmentation. Segmentation results were merged across tiles, and empty polygon geometries were filtered prior to import. Final segmentation results were imported into Xenium - compatible bundle format using XeniumRanger import-segmentation (v3.1.1.0) with spatial coordinates in microns. All processing was orchestrated using Nextflow (v24.10.0).

#### Proseg

Probabilistic transcript-to-cell assignment was performed using Proseg (v2.0.3) in Xenium mode (--xenium). Proseg implements a cellular Potts model that iteratively simulates voxel-level cell boundaries to fit observed transcript distributions, generating cell polygons and probabilistic transcript assignments. The algorithm was run with default parameters on each tissue punch independently, using 4 CPU threads per sample. Proseg outputs—including cell polygons (GeoJSON), transcript-to-cell assignments, expected count matrices, and cell metadata—were converted to Baysor-compatible format using the proseg-to-baysor utility and subsequently imported into Xenium-compatible bundle format using XeniumRanger import-segmentation (v3.1.1.0) with spatial coordinates in microns and 128 GB memory allocation. Proseg was executed within the khersameesh24/proseg:2.0.0 Docker container, orchestrated via a Nextflow pipeline (v24.10.0).

#### Segger

Graph neural network-based cell segmentation was performed using Segger (v0.1.0), which constructs heterogeneous graphs of transcripts and nuclei to assign transcripts to cells. The pipeline consisted of three stages: dataset creation, model training, and prediction. During dataset creation, transcript and boundary node graphs were constructed with k-nearest neighbor parameters of k=3 and maximum distances of 5 µm for transcript nodes and 50 µm for boundary nodes. Data were tiled into 200 × 200 pixel tiles, with tiles smaller than 20 pixels in either dimension excluded to prevent computational failures at tissue edges. A Segger model was trained for up to 100 epochs with a batch size of 1 on a single GPU using CUDA 12.1. Prediction was performed using a k-d tree nearest neighbor method with connected component analysis enabled. Post-prediction filtering retained cells with areas between 20 and 500 µm². A patched Docker image (segger_dev:cuda121-fixed) was used to address a known Qhull precision error (GitHub issue #107) affecting cells with tightly clustered transcripts; the patch catches degenerate geometry cases per-cell and skips them with a warning rather than terminating the pipeline. The Segger pipeline was orchestrated using Nextflow with GPU -enabled Docker containers (8 CPUs, 400 GB memory for training and prediction stages).

### CRISP: Co-expression Rejection In Segmentation Purity

First, tissue-specific mutually exclusive marker gene lists are parsed, where each entry specifies a tissue, a cell type, and one or more marker genes. Markers may be single genes (e.g., *Stmn2* for neurons) or compound markers consisting of multiple genes joined with “+” (e.g., *Cd8a + Ptprc + Cd53 + Itgax* for immune cells). For compound markers, expression values of the constituent genes are summed prior to thresholding, increasing sensitivity for cell types whose individual marker genes are expressed at low levels in spatial data.

Second, for each cell in the dataset, the algorithm determines which cell type markers exceed a user-specified detection threshold. The detection threshold (default = 2 counts) defines the minimum expression level at which a gene (or summed compound marker) is considered detected within a cell. For each cell type, a Boolean expression vector is computed across all cells, and these vectors are assembled into a cell type expression matrix (cell types × cells).

Third, the number of distinct cell type markers co-expressed within each cell is counted by summing the columns of the cell type expression matrix. Cells expressing markers from zero cell types are classified as “no_markers” (unassigned). Cells expressing markers from exactly one cell type are classified as “pure.” Cells expressing markers from two or more mutually exclusive cell types are classified as “impure,” indicating potential chimeric cells arising from segmentation errors.

The overall purity score for a tissue is computed as:

*Purity = 1 − (N_impure_ / N_marker-positive_)*

where N_impure_ is the number of cells co-expressing markers from more than one cell type, and N_marker-positive_ is the total number of cells expressing at least one marker. This formulation restricts the denominator to marker-positive cells, ensuring that the purity score reflects segmentation quality among cells with detectable marker expression rather than being diluted by cells outside the marker panel.

In addition to the overall purity score, CRISP performs pairwise co-expression analysis to identify which cell type pairs are most frequently co-expressed. For each impure cell, the specific combination of co-expressed cell types is recorded. These pairwise frequencies are compiled into a symmetric co-expression matrix and ranked to highlight the most common boundary-error patterns, providing diagnostic information about which cell type interfaces are most susceptible to segmentation artifacts in each tissue.

Confidence intervals for purity scores were estimated using an adaptive bootstrap procedure (default: up to 500 iterations at the 95% confidence level). In the adaptive method, bootstrap resampling of cells was performed in batches of 100 iterations, and the procedure terminated early if the running confidence interval stabilized within a tolerance of 0.01 between consecutive batches. For each bootstrap iteration, cells were resampled with replacement and purity was recalculated. Percentile-based confidence intervals were derived from the empirical distribution of bootstrap purity scores, and the standard error was calculated as the standard deviation of the bootstrap distribution. Alternative confidence interval methods are also implemented, including an analytical Wilson score interval for fast approximation and parallelized bootstrap using multiple CPU cores via the future package.

CRISP also implements automated expression-level warnings to guide threshold selection. When marker expression levels vary substantially across markers within a tissue (coefficient of variation > 50%), when the threshold exceeds the mean expression of any marker, or when mean expression is more than five-fold above the threshold, the software generates diagnostic warnings. These checks help users calibrate thresholds to marker expression levels, reducing the risk of artificially inflated or deflated purity estimates.

CRISP was implemented as both an R package (crispRSeg, v0.1.0) and a Python package (crispySeg). The R implementation provides S3 methods for Seurat (v5.0+) and SpatialExperiment objects, enabling direct integration into standard single-cell analysis workflows. Both implementations accept marker definitions as CSV files or data frames and support tissue-specific thresholds via named vectors. The crispRSeg package additionally includes functions for automated marker discovery from reference single-cell RNA-seq datasets (*find_neg_coexp_markers()*), visualization of purity results, and appending cell-level purity annotations to single-cell object metadata. The source code is available at https://github.com/santangelo-lab/CRISP under an MIT license.

### Simulation studies

To validate the accuracy of CRISP purity estimation and characterize the interaction between detection threshold and marker expression level, simulation studies were conducted using synthetic datasets with known ground truth purity levels. All simulations were performed in R using the crispRSeg package and Seurat (v5.0+).

#### Synthetic data generation

For each simulation, a Seurat object containing 1,000 cells and 50 genes (3 marker genes and 47 non-marker genes) was generated. Of these cells, 80% (800 cells) were designated as marker-positive. Three mutually exclusive marker genes (*Actc1, Acta2, Pecam1*) were assigned to three cell types (cardiomyocyte, smooth muscle, and endothelial), mimicking a heart tissue marker panel. True impurity was introduced at controlled rates of 0%, 5%, 10%, 20%, and 30% of marker-positive cells. Pure cells were randomly assigned to express a single marker gene, while impure cells were programmed to co-express two marker genes (*Actc1* and *Pecam1*), simulating chimeric cells arising from segmentation errors.

Marker gene expression values were drawn from a negative binomial distribution parameterized by a mean expression level (*mu*) and a size parameter controlling overdispersion. Three mean expression levels were tested: *mu* = 4, 10, and 50 counts per cell, representing low, moderate, and high marker expression scenarios typical of different spatial transcriptomics platforms and gene panels. Expression noise was introduced at three levels by varying the negative binomial size parameter: low noise (size = 10), medium noise (size = 5), and high noise (size = 2), where lower size values produce greater variance. Background expression was added at a mean level of 1 count across all genes using a negative binomial distribution (size = 2) to simulate ambient RNA contamination.

#### Threshold sensitivity analysis

CRISP was applied to each simulated dataset across a range of detection thresholds. For simulations with mean expression of 4, thresholds of 1 through 10 were tested; for mean expression of 10 and 50, thresholds were extended to match the expression range (up to 50). At each threshold, the per-cell purity classifications from CRISP were compared against the known ground truth labels. Classification performance was evaluated using sensitivity (recall: proportion of truly impure cells correctly identified), specificity (proportion of truly pure cells correctly classified), precision, F1 score (harmonic mean of precision and recall), and overall accuracy. The false positive rate (pure cells incorrectly flagged as impure) and false negative rate (impure cells missed) were also computed.

#### Purity estimation accuracy

To assess the accuracy of CRISP purity estimates, the estimated purity score (fraction of pure cells among marker-positive cells) was compared against the true purity for each combination of impurity rate, expression level, and detection threshold. This analysis characterized how threshold selection relative to marker expression affects the direction and magnitude of estimation bias. Specifically, setting thresholds above the mean marker expression can cause impure cells to escape detection (artificially inflating purity), while setting thresholds substantially below the mean can cause background expression to be counted as signal (artificially deflating purity). These results informed our recommendation that detection thresholds be set to approximately half the expected mean marker expression for optimal accuracy.

All simulations were seeded for reproducibility (base seed = 42). Bootstrap confidence intervals were disabled during simulation studies to isolate the effects of threshold and expression parameters from sampling variability.

### Downsampling analysis

To evaluate the stability and reliability of CRISP purity estimates across varying sample sizes, a downsampling analysis was performed on real spatial transcriptomics data from the tissue microarray benchmark dataset. Four tissues with varying cell numbers and marker prevalence were selected: heart (∼12,500 cells), liver (∼24,000 cells), small intestine (∼27,000 cells), and spleen (∼108,000 cells). Tissue-specific mutually exclusive marker gene lists were used as defined for the benchmarking analysis.

For each tissue, cells were randomly downsampled to a series of absolute sample sizes: 500; 1,000; 2,500; 5,000; and 10,000 cells, with additional sizes of 20,000 (liver), 25,000 (small intestine, spleen), and 50,000 (spleen) where the full dataset was sufficiently large. The full dataset for each tissue was included as the maximum sample size. At each sample size, 20 independent random subsamples were drawn without replacement to assess both the accuracy and precision of purity estimates. Random sampling was performed on all cells within a tissue, preserving the natural proportions of marker-positive and marker-negative cells.

CRISP was applied to each subsample with a uniform detection threshold of 3 counts and bootstrap confidence intervals disabled, yielding a purity score for each of the 20 replicates at each sample size. This design produced a distribution of purity estimates at each sample size, from which the median purity score, standard deviation, standard error (SD / *sqrt*(n_replicates_)), and coefficient of variation (CV = SD / mean × 100%) were calculated. Parallel processing was performed using the future package in R with 8 workers.

To determine minimum recommended sample sizes, acceptability thresholds of standard error < 2% and CV < 5% were applied. These criteria were evaluated at each sample size for each tissue to identify the minimum ce **l** count at which CRISP purity estimates become sufficiently stable for reliable comparisons. The proportion of cells flagged as impure and marker-specific prevalence (fraction of cells expressing each marker) were also tracked across sample sizes to assess the consistency of impurity detection and marker representation at different scales.

All downsampling was performed on pre-processed Seurat objects derived from the 10x multimodal segmentation output. Random seeds were set for reproducibility (base seed = 42, with unique per-replicate offsets). The analysis was implemented in R using Seurat (v5.0+), the crispRSeg package (v0.1.0), and the future/future.apply packages for parallelization.

### Benchmarking metrics

#### TMA data processing

Due to limitations of the number of region selections available through XOA software, TMA slides were run as one single region and a pipeline for separating individual tissue regions was developed and applied downstream. Following segmentation, the Xenium output bundle was split into individual punch-level datasets using a custom Python (v3.11) pipeline (Xen_TMA_pipeline). Each 2 mm punch region was defined by spatial coordinates, and cells and transcripts were assigned to punches by filtering on centroid position (x_centroid, y_centroid for cells; x_location, y_location for transcripts). Only transcripts with a Phred-scaled quality score (Q-score) > 20 were retained for downstream analysis. For each punch, the pipeline produced a self -contained dataset comprising filtered cell-by-gene expression matrices (HDF5 sparse format), cell and nucleus boundary polygons, transcript coordinates, and cropped morphology images, preserving compatibility with 10x Xenium Explorer and XeniumRanger (v3.1; 10x Genomics). Per-punch quality control metrics were calculated during splitting, including total cell counts, median cell area, median transcripts per cell, and the proportion of decoded transcripts assigned to cells. Segmentation outputs from all five algorithms were processed through this pipeline identically, ensuring that all downstream comparisons were performed on spatially matched tissue regions.

### Segmentation metrics

#### Cell count and morphology

Cell counts, median cell area (in um2), and median transcripts per cell were extracted from each algorithm’s segmented output for all ten tissues. To facilitate cross-algorithm comparison, cell counts were normalized to the nuclear segmentation baseline within each tissue, expressed as a fold-change ratio. Transcript capture rate was defined as the percentage of all decoded transcripts (Q-score > 20) within a punch that were assigned to a segmented cell. Pearson correlation coefficients were calculated between median cell area and median transcripts per cell across all tissue-algorithm combinations, with p-values adjusted using the Benjamini-Hochberg method.

#### Clustering analysis

Unsupervised clustering was performed uniformly across all tissue-algorithm combinations using Seurat (v5.4.0) in R (v4.5.0). Cells were filtered to retain those with a minimum of 10 transcripts, 20 unique genes, and a cell area between 10 and 1,000 um2. Expression counts were normalized using an area-aware log-normalization method that scaled transcript counts per 100 um2 of cell area prior to log-transformation, accounting for the substantial variation in cell size across segmentation algorithms. The top 2,000 variable features were identified using Seurat’s variance-stabilizing transformation, and principal component analysis (PCA) was performed on the scaled variable features. The number of principal components retained for downstream analysis was set per tissue based on elbow plot inspection (range: 30-45 PCs; Supplementary Table 1). Cells were clustered using Seurat’s shared nearest neighbor (SNN) graph-based approach with the Louvain algorithm, with clustering resolution set to 0.2 for brain tissue and 0.4 for all other tissues. UMAP embeddings were computed for visualization.

#### Silhouette width

Clustering quality was assessed by computing silhouette width scores using the cluster package (v2.1.8.1) in R. For each tissue-algorithm combination, Euclidean distances were calculated in PCA space using the same number of PCs as used for clustering. To manage computational cost, a random subsample of up to 5,000 cells was drawn when the dataset exceeded this threshold. Mean and median silhouette widths were reported for each algorithm, and per-cluster mean silhouette widths were calculated for the brain case study.

#### Differential expression analysis

Cluster marker genes were identified using Seurat’s FindAllMarkers function with a log2 fold-change threshold of 0.5 and the Wilcoxon rank-sum test. Genes with Bonferroni-adjusted p-values < 0.05 were considered significantly differentially expressed. For each marker gene, the absolute log2 fold-change and the percent expressed difference (the difference in the fraction of cells expressing the gene between the cluster of interest and all other clusters) were recorded. Distributions of these metrics were compared across segmentation algorithms using one-way ANOVA, with Tukey’s HSD post-hoc test to identify pairwise differences.

#### CRISP purity scoring

Segmentation purity was assessed for each tissue-algorithm combination using CRISP (crispRSeg v0.1.0). Tissue-specific mutually exclusive marker gene sets were curated for each of the ten tissues through prior domain knowledge and consultation of existing single-cell RNA-seq literature (Supplementary Table 2). Marker selection followed several criteria: markers were required to be mutually exclusive across cell types and matched to the tissue of origin, expressed in at least 1,000-10,000 cells in the dataset of interest, and exhibit sufficiently high expression levels for reliable detection. Expression levels between marker pairs were matched as closely as possible; when a single gene was insufficiently expressed relative to its partner, compound markers (sums of multiple genes, denoted with “+”) were used to increase the effective expression level. The detection threshold--the minimum transcript count at which a cell is considered to express a given marker--was set consistently within each tissue, ranging from 1 to 3 depending on the average expression of the selected markers in the nuclear-segmented data. This anchoring to the nuclear segmentation baseline ensured that threshold values reflected biological expression levels rather than algorithm-specific differences in transcript capture. CRISP was run on each tissue-algorithm combination using the tissue-matched marker set and threshold.

### Statistical analysis

All statistical analyses were performed in R (v4.5.0). Pearson correlation coefficients were used to assess relationships between continuous metrics (e.g., median cell area and median transcripts per cell) across all tissue-algorithm combinations (n = 60; 10 tissues x 6 segmentation approaches). All correlation tests were two-tailed. Where multiple correlations were tested simultaneously, p-values were adjusted using the Benjamini-Hochberg method. Distributions of differential expression effect sizes (absolute log2 fold-change, percent expressed difference) were compared across segmentation algorithms using one-way ANOVA followed by Tukey’s HSD post-hoc test for pairwise comparisons; significance groupings were determined at p < 0.05. For the downsampling stability analysis, CRISP purity scores were computed at each sample size across 20 independent random subsamples; median and interquartile range (IQR) are reported. Differential expression testing used the Wilcoxon rank-sum test with Bonferroni correction for multiple comparisons. In all box plots, the center line indicates the median; box limits represent the first and third quartiles, and whiskers extend to 1.5x the interquartile range.

## Data Availability

A full table of benchmark metrics across algorithms and tissues is available in Supplementary Table 3. All Xenium spatial transcriptomics data generated in this study have been deposited in GEO under accession number GSE328169.

## Code Availability

The CRISP R package (crispRSeg) and Python package (crispySeg) are available at https://github.com/santangelo-lab/CRISP under an MIT license. The Nextflow pipeline for performing segmentation is available at https://github.com/jrose835/Xen_Segmentation_NextFlow. The Xen_TMA_pipeline code for processing TMA regions is available at https://github.com/jrose835/Xen_TMA_pipeline. All analysis scripts for reproducing figures are available at https://github.com/santangelo-lab/segmentation_benchmark. Code for this project was written using agentic AI tools.

## Acknowledgements

This material is based upon work supported by the Defense Advanced Research Projects Agency (DARPA) under Agreement No. HR00112590020. The views and conclusions contained in this document are those of the authors and should not be interpreted as representing the official policies, either expressed or implied, of the U.S. Government. The authors acknowledge Vaunita Parihar of the Cancer Tissue and Pathology (CTP) core center at Emory University for her guidance on tissue slide scanning.

## Author Contributions

JR, DV, and PJS conceived the study. JR developed the CRISP software, performed all computational analyses, and wrote the manuscript. ER devised the TMA structure, selected regions for TMA creation, performed staining and Xenium protocols. HP, LS, and CP helped create RNA and LNP reagents necessary for the animal study and TMA creation. JA and HP performed the animal experiment. DV and PJS supervised the study and revised the manuscript. All authors read and approved the final manuscript.

## Materials & Correspondence

Correspondence and material requests should be addressed to Philip J Santangelo

## Supplementary Information

**Supplementary Figure 1.**
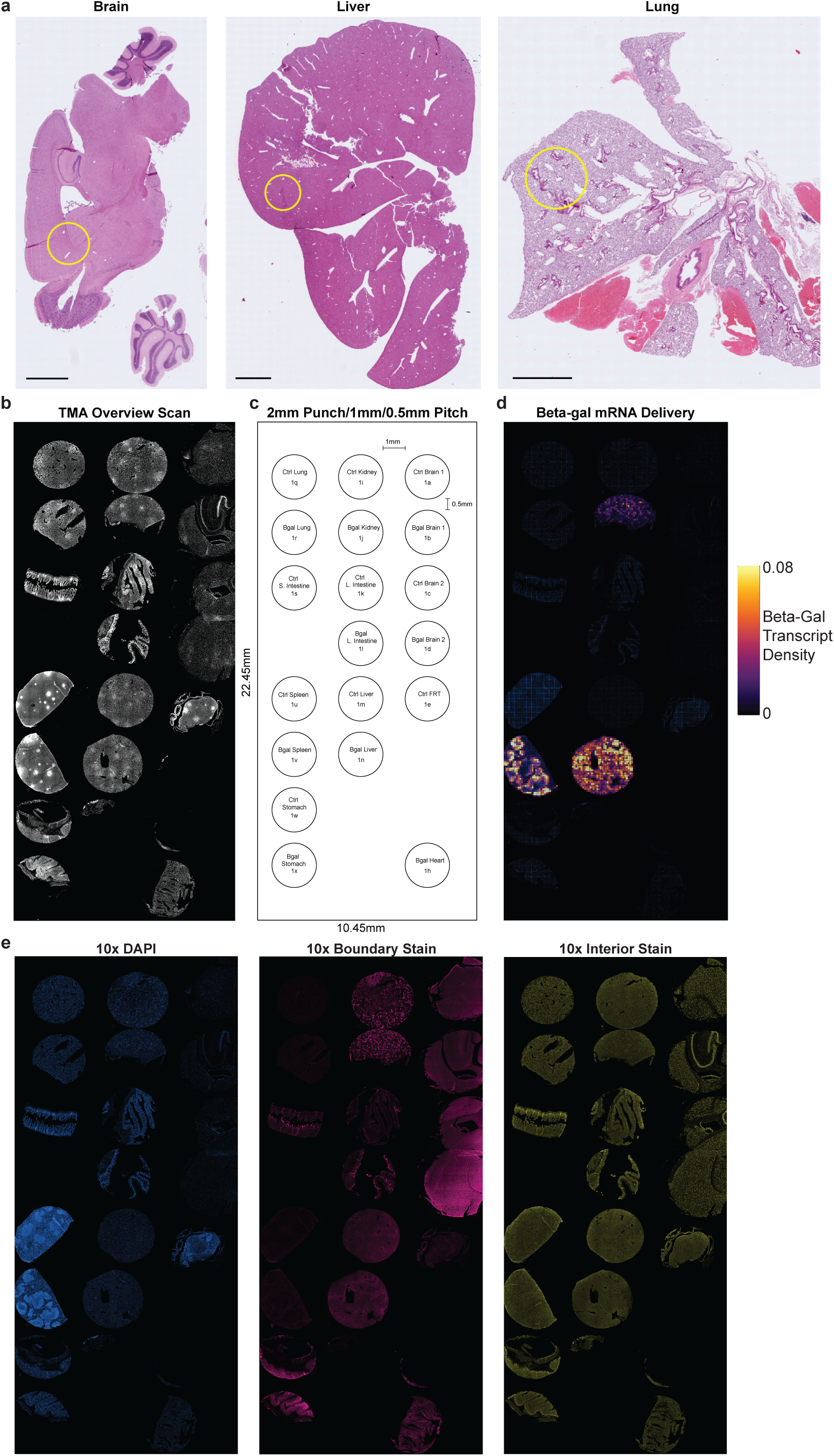
Tissue micro array construction. **a.** Representative H&E-stained whole tissue sections from brain, liver, and lung, with yellow circles indicating the 2 mm diameter regions selected for TMA punch extraction. Scale bars represent 2mm. **b.** Autofluorescence overview scan of the complete TMA block, showing all tissue punches after array construction. **c.** Schematic key of the TMA layout, indicating tissue identity, treatment condition (Ctrl = vehicle control; Bgal = β-galactosidase mRNA LNP-treated), and position for each 2 mm punch. Punches were arrayed at 1 mm center-to-center spacing with 0.5 mm edge-to-edge pitch. **d.** Spatial density map of β-galactosidase mRNA probe binding across the TMA, visualizing the distribution of transcript cargo delivered by lipid nanoparticles to treated animals. **e.** Fluorescence images of the full TMA showing the three staining channels used by the 10x Xenium multimodal segmentation kit: DAPI nuclear stain (left), boundary stain targeting cell surface markers ATP1A1, E-Cadherin, and CD45 (center), and interior stain targeting 18S rRNA (right).

**Supplementary Figure 2.**
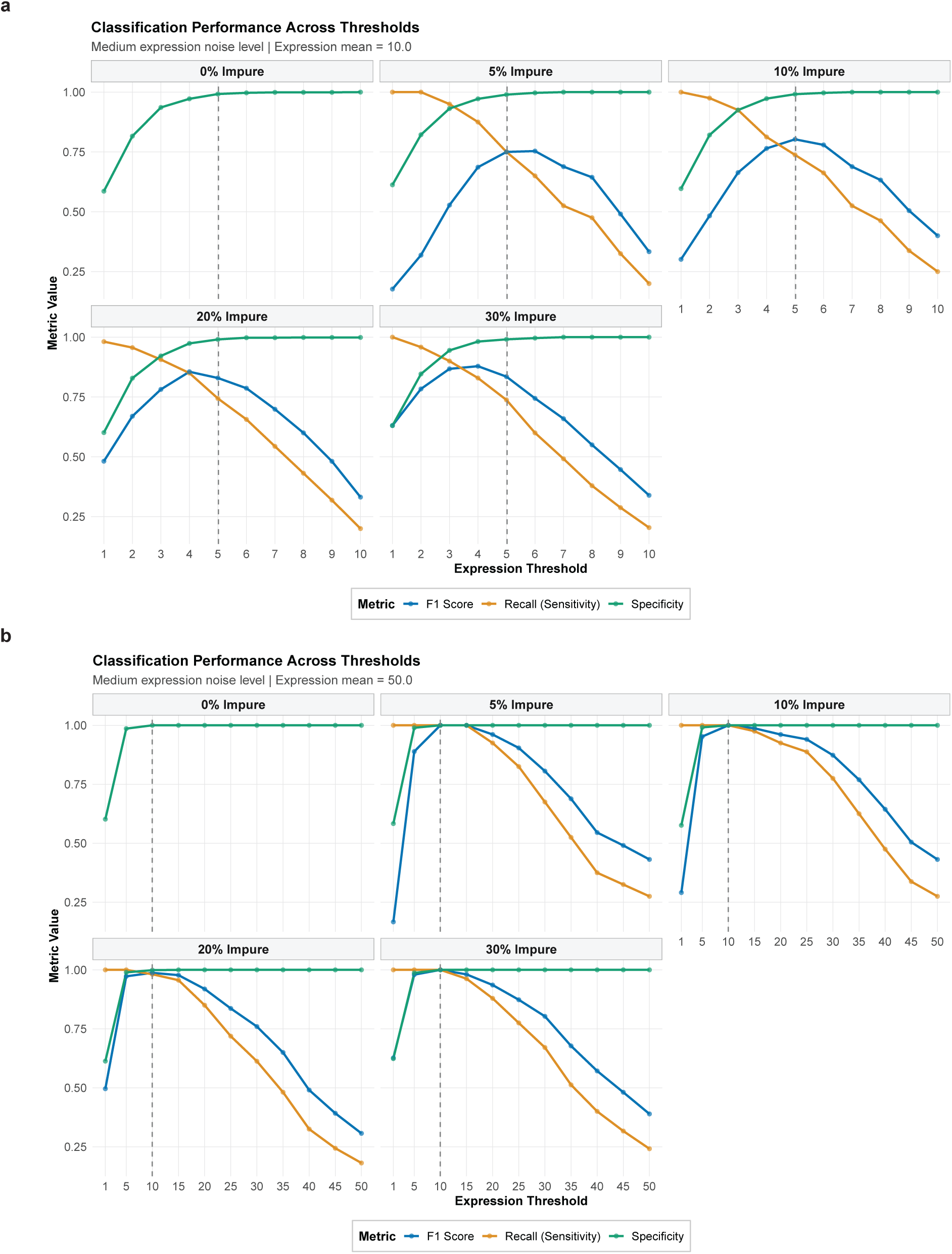
CRISP classification performance at moderate and high marker expression levels. **a.** Classification performance of CRISP across detection thresholds (1–10) at varying levels of simulated true impurity (0%, 5%, 10%, 20%, and 30%) for synthetic datasets with mean marker expression of 10 counts per cell and medium expression noise (negative binomial size = 5). F1 score (blue), recall/sensitivity (orange), and specificity (green) are plotted for each threshold value. **b.** Same analysis as (a) but for simulated mean marker expression of 50 counts per cell, with detection thresholds tested across an extended range (1–50). Expression values drawn from negative binomial distributions with background noise at mean = 1 count.

**Supplementary Figure 3.**
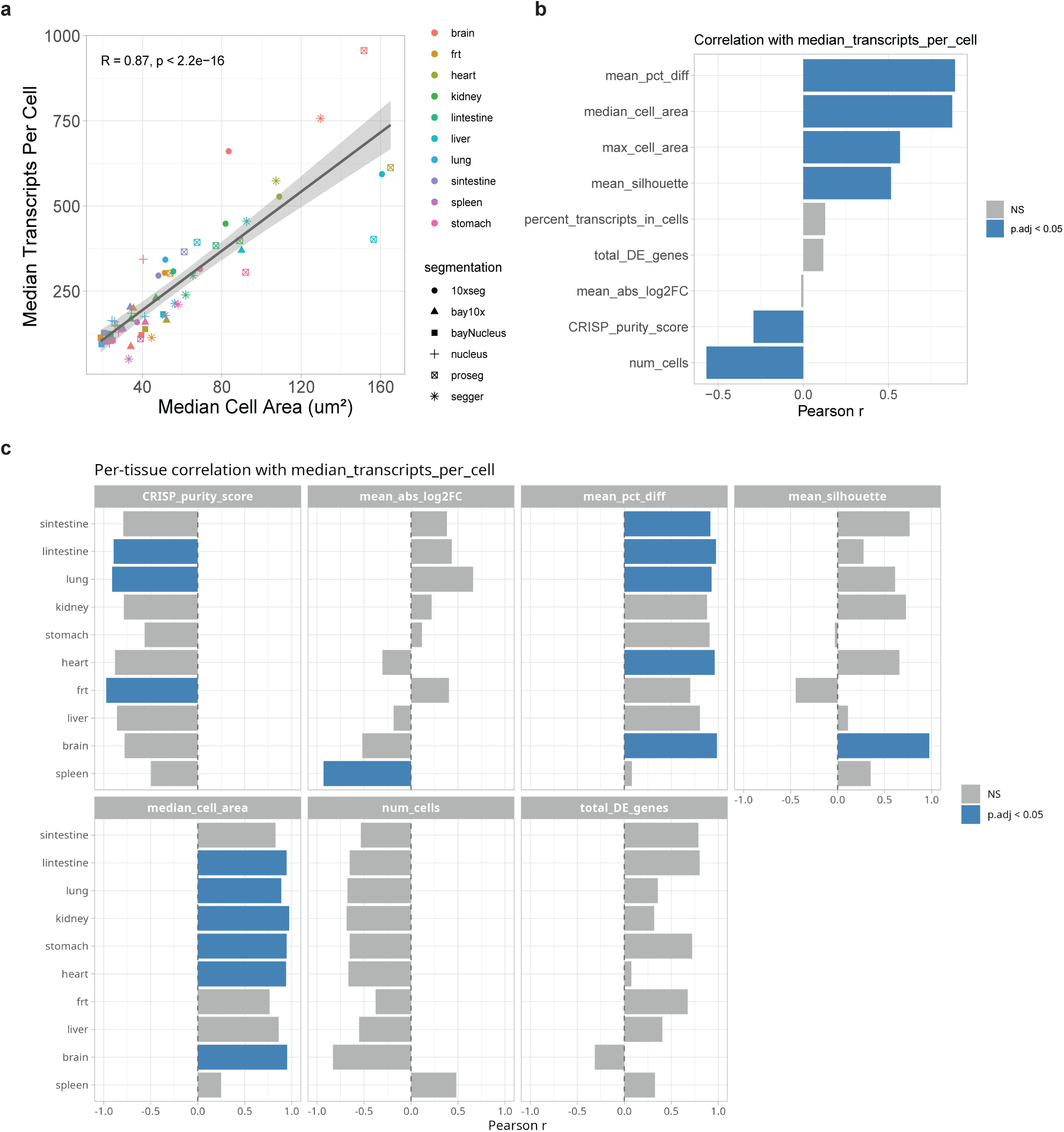
Cell morphology, transcript capture, and downstream metrics are correlated across tissue–algorithm combinations. **a.** Scatter plot of median cell area (µm²) versus median transcripts per cell across all tissue–algorithm combinations (n = 60; 10 tissues × 6 segmentation approaches). Points are colored by tissue and shaped by segmentation algorithm. A strong positive Pearson correlation is observed (R = 0.87, p < 2.2 × 10⁻¹⁶). **b.** Bar plot of Pearson correlation coefficients between median transcripts per cell and each benchmarking metric across all tissue–algorithm combinations. Blue bars indicate statistically significant correlations after Benjamini–Hochberg correction (adjusted p < 0.05); grey bars indicate non-significant correlations. **c.** Tissue-stratified analysis of the correlations shown in (b).

**Supplementary Figure 4.**
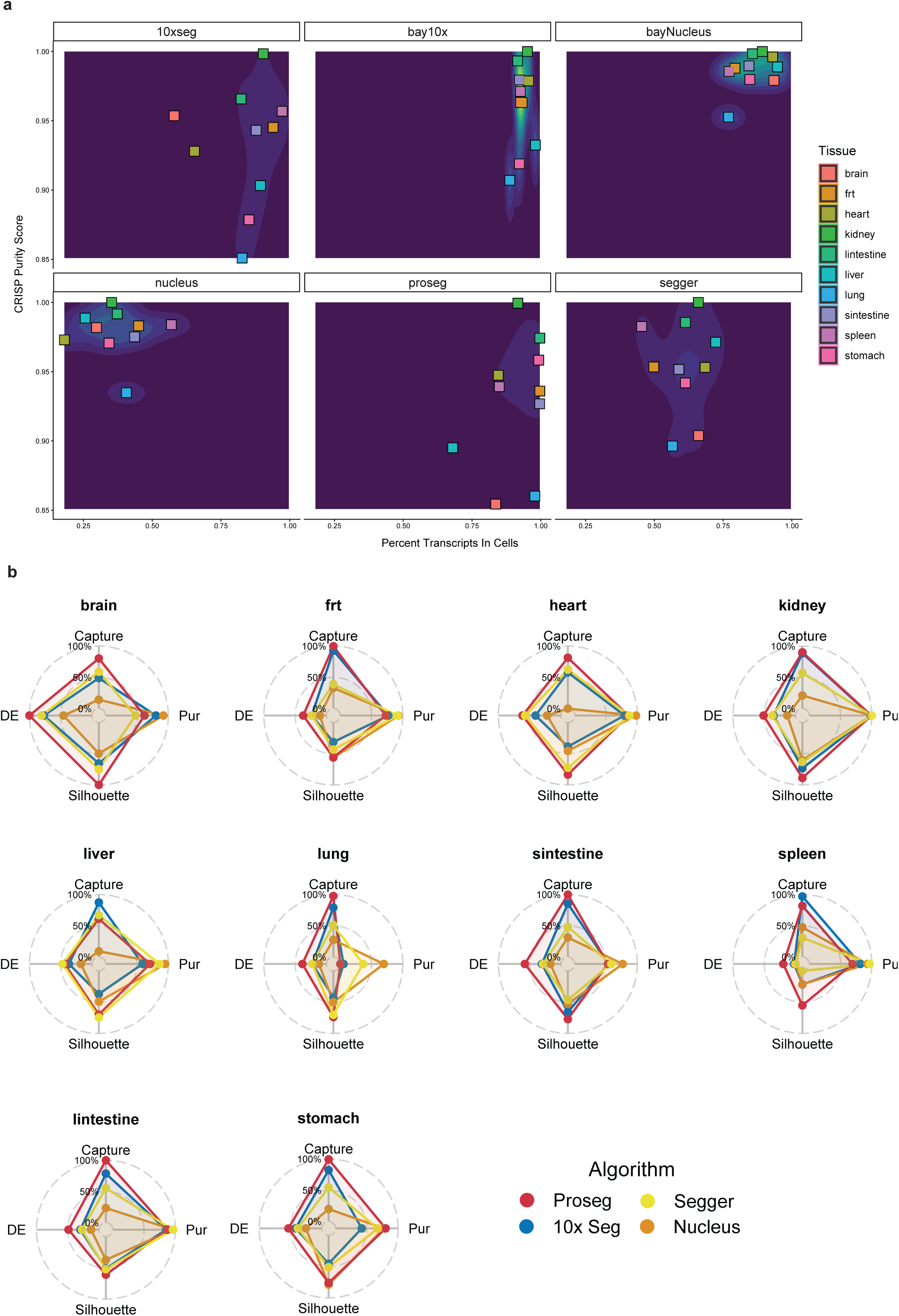
Per-algorithm capture–purity relationships and multi-metric radar profiles across tissues. **a.** Scatter plots of CRISP purity score versus percent of transcripts assigned to cells, faceted by segmentation algorithm, with each point representing a single tissue colored by tissue identity. **b.** Radar plots summarizing algorithm performance across four metric categories for each of the ten tissues: differential expression (DE, combining mean absolute log2 fold-change and mean percent difference), transcript capture (percent transcripts in cells), clustering quality (mean silhouette width), and purity (CRISP purity score). Axes represent min–max scaled values within each tissue (0–100%), enabling direct comparison of relative algorithm performance. Four algorithms are shown for clarity: 10x segmentation kit (blue), nuclear segmentation (orange), Proseg (red), and Segger (yellow).

**Supplementary Figure 5.**
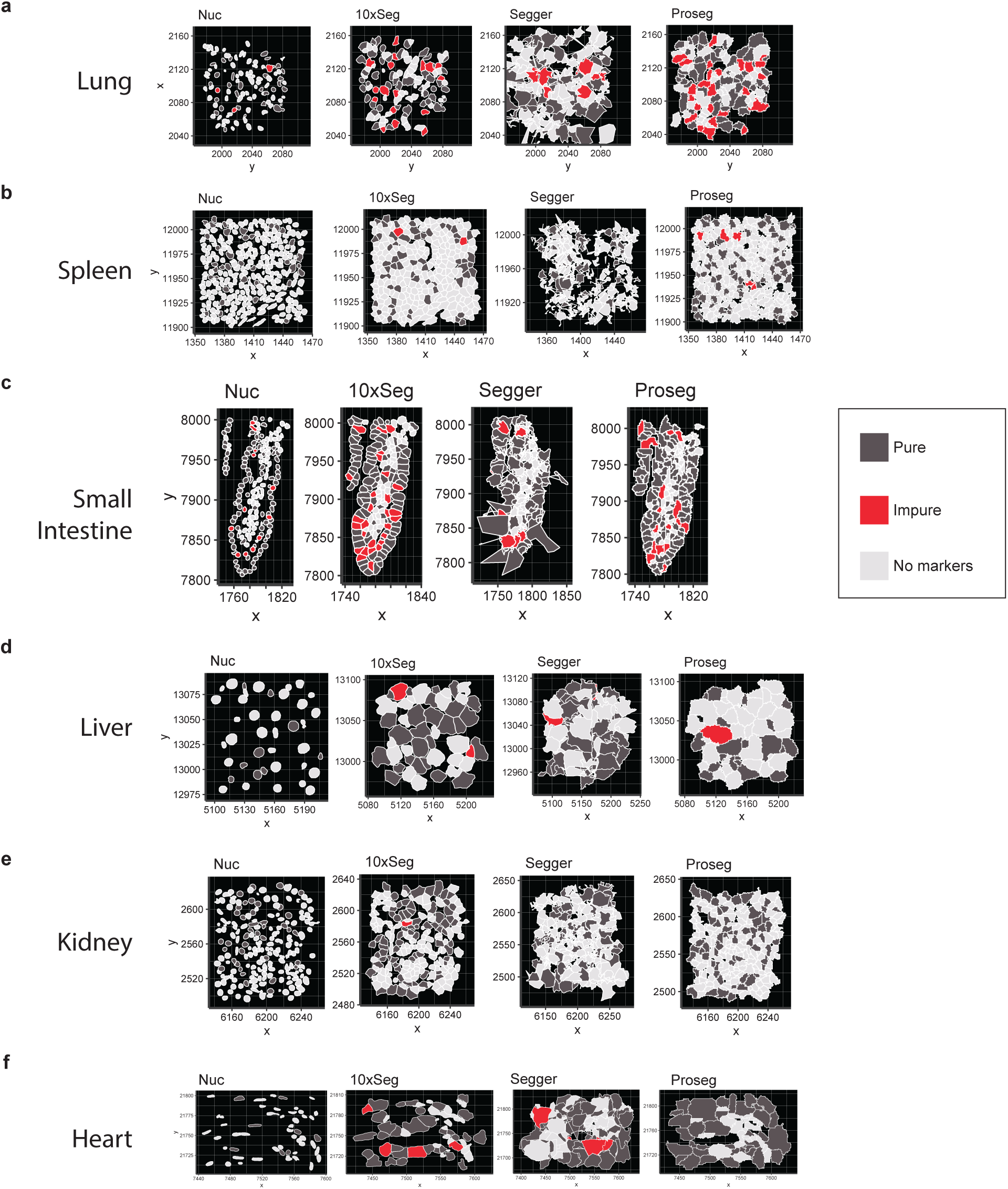
Spatial visualization of cell boundaries and CRISP purity classifications across tissues and algorithms. Spatial maps of CRISP purity classifications overlaid on cell boundary polygons for representative regions from six tissues as indicated, comparing nuclear segmentation (Nuc), 10x segmentation kit (10xSeg), Segger, and Proseg. Cells are colored by purity status: pure (expressing markers from exactly one cell type; dark grey), impure (co-expressing mutually exclusive markers from two or more cell types, indicating potential chimeric cells; red), or no markers (expression below detection threshold for all markers in the panel; light gray). Cell boundaries are drawn as polygons output by each algorithm, displayed at the same spatial scale within each tissue.

